# The Spatiotemporal Proteome Landscape of Aging: Structural determinants of age-sensitive proteome remodeling

**DOI:** 10.64898/2026.02.26.708310

**Authors:** Seungmin Yoo, Lingraj Vannur, Liying Li, Christopher Young, Qingqing Liu, Zhihui T. Wen, Ying Zhang, Laurence Florens, Kausik Si, Junming Zhuang, Fan Zheng, Chuankai Zhou

**Affiliations:** Buck Institute for Research on Aging, 8001 Redwood Blvd., Novato, CA, 94945, USA; Stowers Institute for Medical Research, 1000 East 50th Street, Kansas City, MO 64110, USA

**Keywords:** aging, protein localization, protein concentration, protein structure

## Abstract

Aging is marked by a decline in cellular functions accompanied by widespread changes in mRNA and protein abundance, yet whether aging broadly remodels subcellular protein localization and concentration—and why some proteins change while others remain stable—remains unclear. This gap matters because cellular function depends not only on expression levels but also on correct spatial organization. Using yeast replicative aging as a model, we built a robotic pipeline to enrich old cells from 5,661 strains, acquired 90 million single-cell 3D images, and applied machine learning to map proteome-wide changes in localization, concentration, and aggregation throughout aging. This age-resolved single-cell atlas uncovers widespread proteome remodeling and rewiring of protein interaction networks. Moreover, structural analysis reveals biophysical determinants of age-sensitive proteome remodeling across ages and species. Together, these results reveal a structure-encoded intrinsic principle underlying spatial proteome breakdown during aging and provide a resource to dissect mechanistic links among aging hallmarks.

## Introduction

Eukaryotic cells compartmentalize their contents and functions into various membrane-bound organelles, creating specialized microenvironments with precisely regulated concentrations of proteins and metabolites that support metabolism and cellular functions within each organelle^1^. With age, this spatial organization and functional integrity progressively deteriorate, giving rise to conserved hallmarks such as genomic instability, proteostasis decline, mitochondrial dysfunction, and disrupted inter-organelle communication^2–4^. Although these hallmarks are well-described outcomes of aging, the molecular processes that drive their emergence remain poorly understood^5^. A central gap is how the proteome—the complete set of cellular proteins—progressively changes over time in localization, concentration, and interaction states, and how these proteome-wide reorganizations contribute to the decline of cellular functions during aging.

Over the past decade, transcriptomic and proteomic approaches have transformed our understanding of age-associated gene expression changes^6^. Recent single-cell transcriptomic atlases have further resolved cell-to-cell heterogeneity and aging trajectories, yet the temporal dynamics of proteome remodeling at single-cell resolution remain largely unexplored. This gap limits mechanistic interpretation of how transcriptional programs translate into functional decline, particularly given the well-documented age-dependent uncoupling between mRNA and protein abundance^7–9^. Moreover, most omics studies primarily quantify mRNA and protein abundance, leaving unresolved how the spatiotemporal localizations and local concentrations of proteins change across organelles during aging. This gap is critical because proteins function not only through coordinated expression, but also through precise control of their spatial distribution and local concentration, formation of macromolecular complexes, and dynamic interactions with organelles and other biomolecules^10–12^. Accordingly, subcellular localization studies of individual proteins have frequently been central to uncovering mechanisms of aging, yet we still lack a comprehensive, age-resolved atlas of proteome localization and concentration. This limitation primarily stems from technical challenges associated with systematic measurements of the subcellular localizations and concentrations of proteins, particularly in replicative aged cells which are rare and fragile in model organisms^13^. Previous studies from different laboratories have investigated age-associated expression and localization of individual proteins in hypothesis-driven contexts, but variability in methods and genetic background has made it challenging to systematically synthesize aging mechanisms across multiple organelles^14^. Thus, a comprehensive, age-resolved atlas of proteome localization and concentration changes in eukaryotic cells remains an urgent gap to be filled.

To address these gaps, we performed a comprehensive, proteome-wide, single-cell analysis of protein dynamics during replicative aging in budding yeast—a genetically tractable system that has long served as a powerful model for uncovering conserved mechanisms of aging^14,15^. As replicative aged cells are rare, we developed a high-throughput robotic pipeline to enrich aged cells across 5, 661 isogenic strains, each expressing a single endogenously tagged protein, and to acquire over 90 million three-dimensional fluorescence images of individual young and aging cells. Using convolutional neural networks trained to recognize subcellular compartments and morphological features, we systematically quantified localizations, concentrations, interactions, and aggregation states for 5,661 proteins, representing 94% of the entire yeast proteome, across the replicative lifespan at single-cell and single-age resolution.

This single-cell, single-age–resolved proteome atlas reveals hundreds of previously unrecognized, age-dependent protein changes across organelles—many of which are not accessible to bulk proteomics because they manifest as subcellular relocalization, aggregation, and remodeling of protein concentration and interaction networks, and are further obscured by pronounced cell-to-cell heterogeneity during aging. By integrating imaging readouts with protein structural information, we find that intrinsic structural features of proteins—such as surface chemistry and compactness—predict their susceptibility to age-associated disorganization across ages and species, highlighting a biophysical basis for proteome disorganization during aging. This spatiotemporal proteomic map provides a resource to illuminate how aging cells reorganize their proteome and to dissect the processes that give rise to distinct hallmarks of aging, offering a framework for linking molecular-level changes to systems-level aging phenotypes.

## Results

### High-throughput robotic pipeline to enrich and image replicative aged cells

To explore the age-dependent spatiotemporal changes in protein expression, localization, and aggregation, we implemented a single-cell imaging pipeline to examine each of the 5,661 proteins that seamlessly C-terminally tagged with mNeonGreen (mNG) at their endogenous loci^16^. We selected imaging rather than mass spectrometry because enriched populations of replicative aged cells invariably consist of mixed ages^8^, which mass spectrometry cannot distinguish. Imaging provides both single-cell and single-age resolution, enabling detection not only of abundance changes but also of subcellular localization and local concentration changes. The seamless mNG library was generated using scarless endogenous tagging that removes the selection marker and preserves the native 3′UTR, minimizing perturbations to endogenous post-transcriptional regulation^16,17^. The mNG collection has been rigorously validated through comparison with multiple orthogonal yeast proteome datasets, including microscopy of the GFP collection, mass spectrometry–based proteomics, and immunoblot-based measurements of protein abundance^16^. Consistently, we confirmed that protein abundances measured in young cells from our mNG collection closely match prior microscopy- and mass spectrometry–based datasets (Figure S1A–B).

A major technical challenge in systematically characterizing proteome changes during aging is that replicative aged cells are rare and must be enriched before analysis^13^. Existing enrichment methods rely on manual, labor-intensive protocols that are costly and impractical to scale across the full library of 5,661 strains. To overcome this limitation, we adapted the enrichment protocol and developed an automated, high-throughput pipeline using a Tecan Fluent liquid-handling robot, enabling reproducible old-cell enrichment at scale (Figure 1A, S1C). Enriched replicative aged cells were stained with calcofluor white (CW) to delineate the cellular boundaries and Wheat germ agglutinin (WGA) to mark bud scars for age quantification. Cells were then imaged by high-throughput spinning-disc confocal microscopy, acquiring 3D z-stacks in the mNG, CW, and WGA channels. In total, we collected 467,456 multidimensional image stacks comprising more than 79.4 million live cells spanning a wide range of replicative ages, along with 10.7 million young cells from overnight culture as controls. Individual cells were subsequently segmented using CellPose for downstream analysis (Figure 1B).

**Figure 1.**
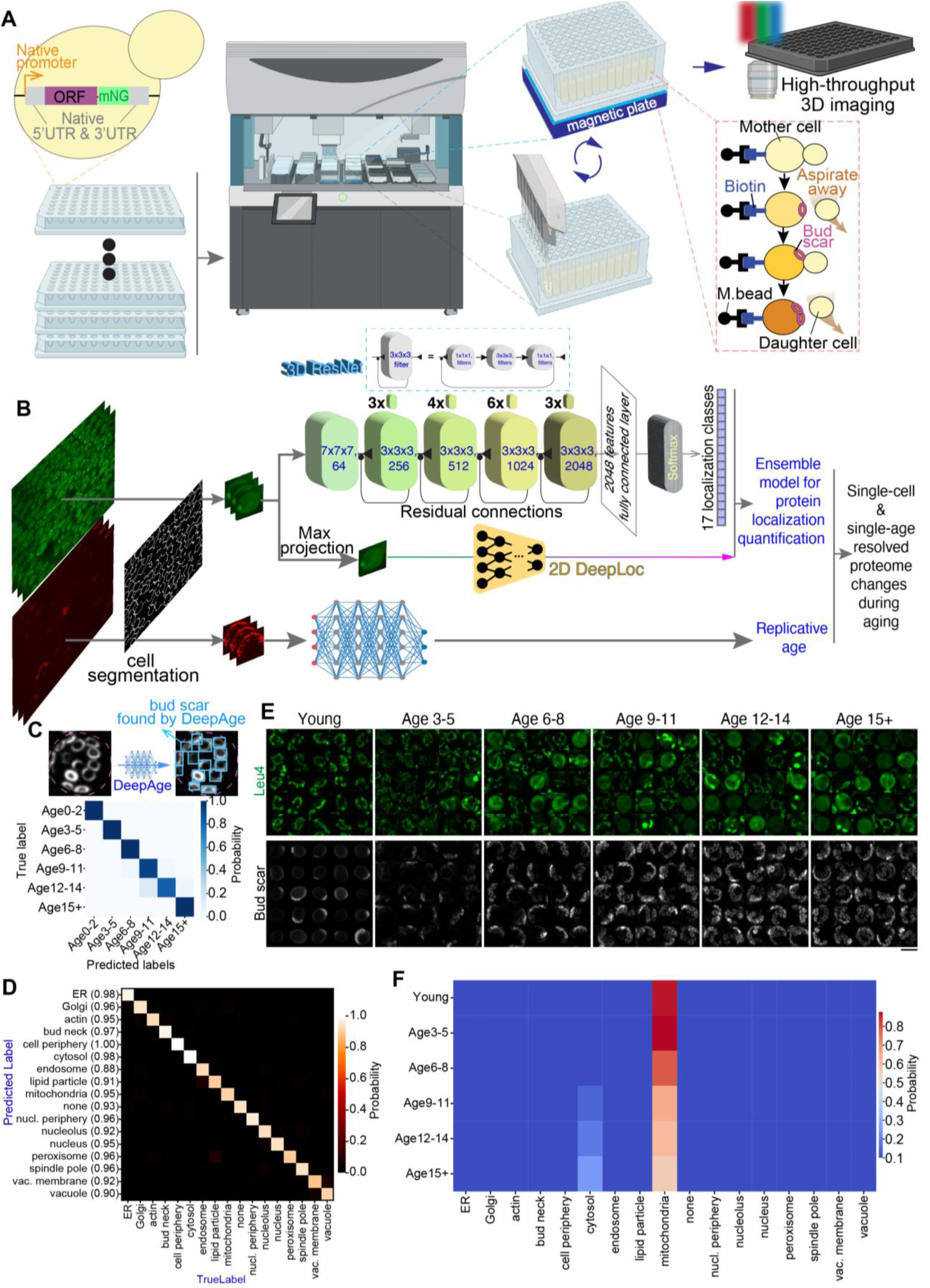
A high-throughput pipeline for collecting aged cells and spatiotemporal analysis of proteome. (**A**) Overview of the yeast library, automated old cell enrichment, and high-throughput 3D imaging pipeline. A 96-deepwell plate was transferred to a magnetic plate every two hours to retain aging mother cells while newborn daughter cells were aspirated and removed by the robot. M. bead, magnetic beads. (**B**) Imaging analysis pipeline for detecting subcellular localization of proteins and cellular age. (**C**) Confusion matrix for the performance of DeepAge in detecting cellular ages. The fraction of cells classified for each age class is shown. (**D**) Confusion matrix for the performance of ensemble model in detecting subcellular localization of proteins. Numbers in brackets on the y axis are the F1 scores (harmonic mean of precision and recall) for each class. Overall F1 score: 0.945. (**E**, **F**) Representative images and localization score of Leu4 as an example of age-associated change of protein localization quantified by the ensemble model. Scale bar: 5μm.

To determine the replicative age of individual cells, we developed a supervised neural network based on the YOLO architecture, which we call DeepAge, to count bud scars in 3D images (Figure 1B). We manually annotated bud scars in 573 single cells, using 77% of these labeled cells to train DeepAge and the remaining 23% to evaluate performance. The resulting model achieved 93% accuracy (Figure 1C). We then applied DeepAge to 79.4 million single cells and assigned each cell to an age bin (0–2, 3–5, 6–8, 9–11, 12–14, and 15+). Because the 0–2 age group includes both young mothers and newly detached daughters from aged mothers generated during sample preparation and live imaging, we used cells from overnight cultures as the young-cell control.

To quantify dynamic protein localization across age-stratified cells, we first retrained DeepLoc, a neural network originally developed for 2D microscopy with additional fluorescent reference markers^12^, using max-projection images from the mNG z-stacks to obtain probabilistic localization calls^18^. Although DeepLoc performed well overall (accuracy ∼89%), several localization classes remained difficult to distinguish even after optimized transfer learning (Figure S1D). This limitation is expected because multiple organelles—including endosomes, lipid droplets, peroxisomes, and Golgi—share similar punctate morphologies and are dispersed throughout the cytoplasm in 2D images (Figure S1E).

We reasoned that leveraging the full 3D information of organelle distribution in our dataset would improve compartment resolution even without additional reference markers. To this end, we developed a new convolutional neural network based on a 3D ResNet framework^19^. In this architecture, fluorescent z-stacks are processed through 3D convolutional blocks, enabling volumetric filters to extract features jointly across the x, y, and z dimensions (Figure 1B). This allowed the model to learn subcellular compartment/organelle-specific 3D shape and texture signatures while retaining spatial invariance. Residual connections are incorporated to bypass one or more layers, facilitating efficient gradient propagation and enabling deeper model training without performance degradation. The final network comprises four residual stages (50 layers total), with batch normalization after each convolution to stabilize training, followed by global average pooling and a fully connected classifier that predicts 17 subcellular localization categories (Figure 1B). In total, the model contains over 46.2 million trainable parameters and achieves robust performance in extracting and classifying features from 3D single-cell images (see STAR Methods).

We trained the 3D ResNet model on ∼29,820 manually annotated z-stacks spanning 17 localization classes (16 subcellular compartments and a ‘none’ class representing no expression), using the same dataset framework as our DeepLoc training. The 3D ResNet model outperformed transfer-trained DeepLoc across nearly all classes, achieving 94% mean accuracy and an F1 score of 0.937 without requiring additional fluorescent reference markers that were needed for the 2D DeepLoc model (Figure S1F)^12^. This improvement likely reflects the fact that organelles exhibit compartment-specific 3D shape and texture signatures; by integrating information across z-slices, the model captures volumetric spatial context that is lost in 2D projections, enabling more accurate discrimination of visually similar localization patterns.

Although the 3D ResNet model outperformed 2D DeepLoc in these tests, there were still cases where 2D DeepLoc worked better, likely because the max-projection of images enhanced signal-to-noise contrast for some low signal proteins. To leverage the complementary strengths of both approaches, we constructed an ensemble model that combines 3D-ResNet with transfer-trained DeepLoc, improving performance to 95% mean accuracy and an overall F1 score of 0.945 (Figure 1D; STAR Methods). We then applied this ensemble model to each cell to obtain probability-based localization profiles across 16 subcellular compartments over replicative ages for each protein (Figure 1B; Table S1). For each strain, we pooled single-cell ages and localization probabilities and binned cells by age to compute age-stratified average localization scores and to detect age-associated protein relocalization (an example, Leu4 relocalization from mitochondria to cytosol, is shown in Figure 1E-F).

### Proteome-wide, age-dependent remodeling of subcellular distribution

We next visualized proteome-wide subcellular protein localizations and their age-associated changes. Among the 5,661 proteins examined, 3,866 displayed clear and specific localization patterns in young cells (hereafter referred to as the localization map; Figure 2A). Across replicative ages, a large fraction of the proteome exhibited age-dependent shifts in subcellular distribution, evident as movement on the spatial proteome map (Figure 2A–C; Video S1). In addition to age-associated changes in mean localization scores, we observed increased heterogeneity in localization across cells for each protein, consistent with the stochastic nature of aging (Figure S2A). Collectively, these age-dependent shifts blur compartmental boundaries, transforming the neatly segregated spatial proteome of young cells—where proteins exhibit characteristic distribution morphologies—into a more overlapping, cross-compartment organization in aged cells (Figure 2A versus 2B).

**Figure 2.**
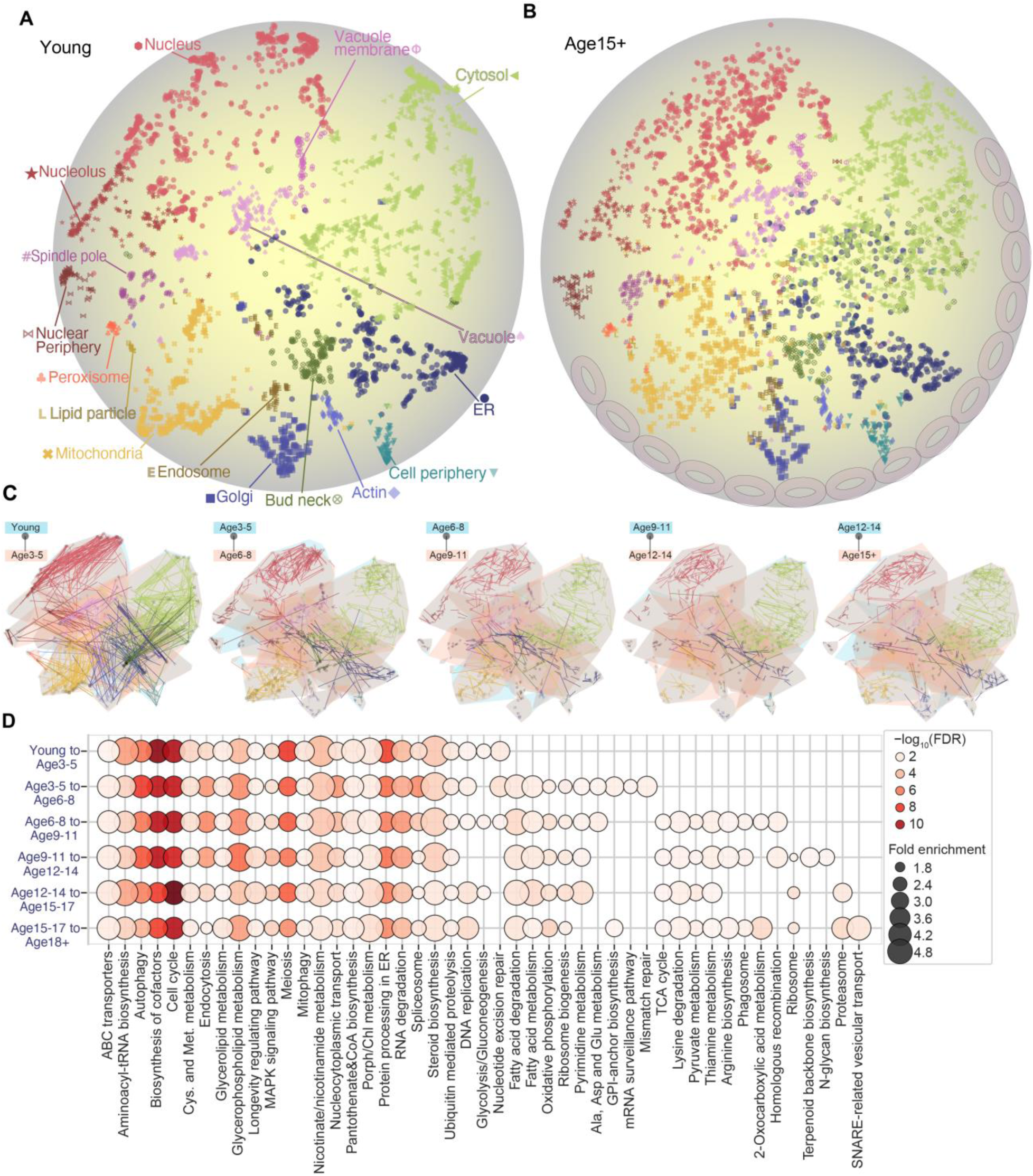
The landscape of spatiotemporal proteome dynamics during aging. (**A**, **B)** The localization map of yeast proteome in young (A) and aged cells of 15-plus generation old (B). The quantified localization scores of 3, 866 strains with detectable expression of proteins from the ensemble model were plotted in tSNE space and normalized for comparison between different ages. Each dot represents a protein. **(C)** Global view of age-associated shifts in protein localization across age groups. For each protein, positions at different ages are connected by a line colored according to its compartment assignment in (A). Organelle boundaries were defined by convex hulls and colored by age group. **(D)** KEGG pathway enrichment for proteins showing substantial (>1.5-fold) localization changes across age groups.

Proteome-wide spatial profiling across replicative age showed that most proteins change within their original compartments rather than relocalizing to entirely different organelles (Figure 2C). These within-compartment shifts—reflected as age-dependent changes in localization morphology—likely captured underlying molecular and metabolic reorganization and protein aggregation during aging. Notably, localization remodeling was already prominent in early aging (Figure 2C), consistent with prior work identifying a metabolic inflection associated with vacuolar de-acidification during the first 3–5 divisions^20,21^. Proteins that underwent these age-dependent morphological changes were enriched for distinct pathways that emerged at different stages (Figure 2D, Table S2). For example, proteins exhibiting early shifts (3–5 divisions) were enriched for autophagy and metabolic pathways involving cofactors, glycerophospholipids, nicotinamide, and sulfur-containing amino acids (e.g., cysteine), whereas proteins exhibiting later stages shifts were enriched for proteasome-related functions and SNARE-mediated vesicular transport. Together, these stage-specific enrichments mirror known early changes in vacuolar acidity and sulfur/amino-acid metabolism during replicative aging^20,21^.

We noticed that proteins were not uniformly distributed within each cellular compartment in the localization map; instead, they formed local clusters, suggesting that proximity in the map captures additional relationships beyond organelle identity. To relate these spatial patterns to biological processes, we applied Spatial Analysis of Functional Enrichment (SAFE), which annotates dense regions of a network with enriched functional attributes^22^. In young cells, SAFE identified 655 significantly enriched Gene Ontology (GO) biological process terms that grouped into 42 network regions (Figure S2B, Table S2). Indeed, proteins that clustered within the same organelle were functionally related and enriched for coherent GO terms, providing a structured functional overlay on the localization map (Figure S2B). These results are consistent with the idea that quantitative localization scores from our ensemble model encode compartment identity while also preserving sub-organelle-level functional organization. With age, this sub-organelle-level functional clustering became less distinct, consistent with intra-compartmental redistribution of proteins (Figure S2C). For example, many ribosome-assembly and rRNA-processing factors became dispersed on the localization map and increasingly overlapped with nuclear proteins (Figure S2D-E).

### Proteins relocalization between different cellular compartments during aging

Although most proteins shift within their original compartments, we also detected a distinct set of 215 proteins that exhibited age-associated cross-compartment relocalization events, in which these proteins moved from their canonical compartment in young cells to a different site in aged cells (Table S3). The nucleus was the most frequent destination (Figure 3A), with proteins from multiple compartments disproportionately relocalizing into the nucleus. Beyond the nucleus, substantial relocalization was also observed to and from the cytosol, nucleolus, mitochondria, and endoplasmic reticulum (ER) during aging (Figure 3A, Table S3).

**Figure 3.**
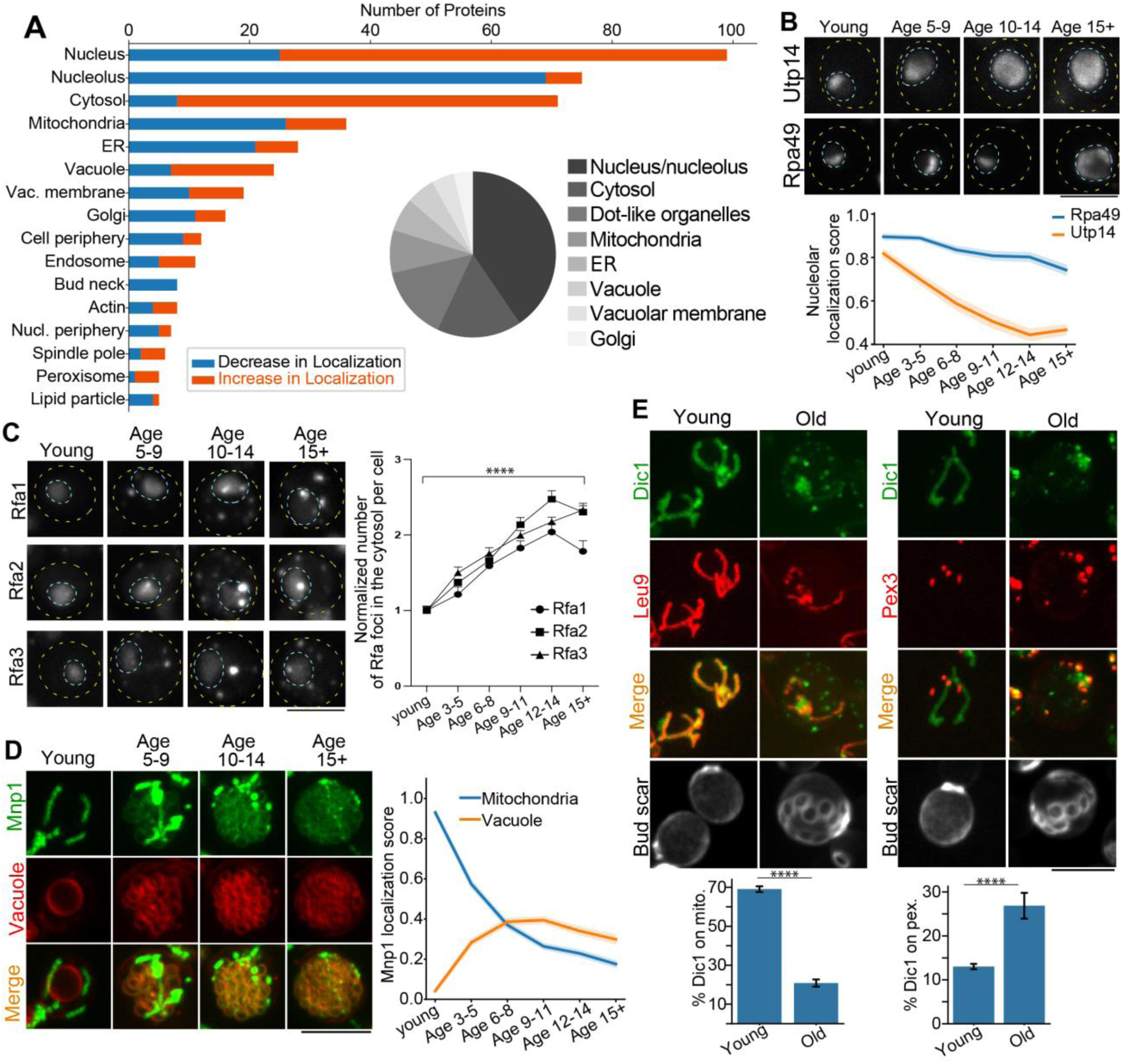
Proteins with age-associated cross-compartment localization changes. (**A**) Number of proteins with inter-compartment localization changes for each organelle. The inset pie chart shows the original localization of these proteins with cross-compartment localization change. (**B**) Representative cells and the nucleolar localization scores of Utp14 and Rpa49 at different ages. (**C**) Representative cells and the quantification of cytosolic foci of Rfa1/2/3 proteins at different ages. **(D**, **E)** Representative cells and the co-localization quantification of Mnp1 (D) and Dic1 (E) at different ages. Shown are Mean and SEM. Scale bar: 5μm. See Source Data table for number of cells quantified.

<u>Nucleolus-to-nucleoplasm redistribution</u>. A prominent class of changes involved nucleolar proteins (e.g., Utp14) shifting into the nucleoplasm, consistent with age-associated perturbation of ribosome biogenesis and nucleolar organization (Figure 3A, 3B, S2E). We also observed gradual dispersion of Rpa49 (a Pol I subunit involved in rDNA transcription) toward the nuclear periphery at advanced replicative ages (Figure 3B). This pattern is consistent with prior reports that Extrachromosomal rDNA circles (ERCs) become abundant in old mother cells and can associate with the nuclear periphery and nuclear pore complexes (NPCs) ^23,24^. However, relocalization of some ribosome biogenesis factors occurred earlier: for example, Utp14, a component of the small subunit (SSU) processome, shifted substantially earlier than Rpa49 and accumulated primarily within the nucleoplasm rather than at the nuclear periphery (Figure 3B). Together, these observations suggest that nucleolus-to-nucleoplasm redistribution of ribosome biogenesis proteins is unlikely to be solely a consequence of ERC accumulation.

<u>Nuclear periphery-to-nucleoplasm shift of an NPC factor</u>. We also observed a pronounced age-associated redistribution of Nup2 from the nuclear periphery into the nucleoplasm (Figure S3A). Because Nup2 is an FG-repeat nucleoporin that interacts with nuclear transport receptors and promotes trafficking through the NPC, its release into the nucleoplasm may contribute to age-dependent defects in nucleo-cytoplasmic transport^23,25^. Notably, this behavior appears selective: most other NPC components remained confined to the nuclear periphery during aging and showed much less relocalization, although several showed age-associated reductions in expression, consistent with previous reports (Figure S3B, S3C) ^23,25^.

<u>Nuclear-to-cytoplasmic shift of genome-maintenance factors</u>. Another notable class of nuclear protein relocalization involves the nuclear-to-cytoplasmic shift. The conserved ssDNA-binding RPA complex (Rfa1/2/3) formed cytosolic foci in addition to nuclear foci (Figure 3C), consistent with both nuclear DNA damage and cytosolic accumulation of ssDNA. Cytosolic DNA has been reported in human senescent cells, where it activates innate immune signaling via the cGAS–STING pathway and contributes to the senescence-associated secretory phenotype (SASP)^26^. In aged yeast, cytosolic Rfa1/2/3 foci showed partial colocalization with mitochondria but not other organelles (Figure S3D), suggesting that mtDNA may contribute to cytosolic ssDNA and that age-associated mitochondrial dysfunction may promote mtDNA release into the cytosol, analogous to human cellular senescence^26^.

<u>Relocalization of mitochondrial proteins</u>. Beyond nuclear proteins, we observed prominent relocalization of mitochondrial proteins. For example, Leu4 accumulated in the cytosol (Figure 1E), Mnp1 redistributed to the vacuolar membrane (Figure 3D, S3F), and Dic1 shifted to peroxisomes (Figure 3E). Leu4 normally functions with its paralog Leu9 as a mitochondrial α-isopropylmalate synthase at the entry point of leucine biosynthesis^27,28^. Its age-associated relocalization to the cytosol—while Leu9 remains mitochondrial (Figure S3E)—could physically separate these isoenzymes and partition leucine biosynthesis across compartments^28^, potentially decoupling mitochondrial leucine availability from cytosolic leucine/TORC1 signaling^29–31^. Mnp1 is a mitochondrial matrix large-subunit ribosomal protein (bL12m) and lacks a transmembrane domain^32^, so its association with the vacuolar membrane in aged cells is unexpected. Its selective relocalization (with other mitoribosomal proteins remaining mitochondrial; Figure S3G) suggests an extra-ribosomal role. Given that Mnp1 is essential for mitochondrial respiration^32^, its age-associated redistribution may contribute to declining mitochondrial translation in aged cells^33^. Dic1 is a mitochondrial dicarboxylate carrier that exchanges malate and succinate for inorganic phosphate across the inner mitochondrial membrane^34^. Its relocalization to peroxisomes could both deplete mitochondria of this carrier and establish dicarboxylate–phosphate exchange at the peroxisomal membrane, thereby rewiring malate/succinate flux among peroxisomes, mitochondria, and the cytosol.

<u>Relocalization of ER proteins</u>. We also observed some ER proteins shifting into dot-like structures, including Hfd1, Erg1, Ice2, and Scs2 (Figure S3H). Hfd1 is a fatty aldehyde dehydrogenase involved in ubiquinone and sphingolipid metabolism; loss of HFD1 causes coenzyme Q (CoQ) deficiency^35–37^. Given that CoQ levels decline with age^38^, mislocalization of Hfd1 may impair CoQ biogenesis even when total Hfd1 abundance is maintained. Erg1 is a rate-limiting enzyme in ER-based ergosterol biosynthesis^39^, Ice2 regulates ER membrane biogenesis and lipid utilization^40^, and Scs2 organizes ER–plasma membrane contact sites and recruits FFAT-motif lipid-handling proteins, contributing to phospholipid and phosphoinositide homeostasis^41,42^. Collectively, these relocalization events are consistent with extensive remodeling of ER architecture and lipid metabolism during aging^43,44^.

### The landscape of inter-compartment protein relocalization and connectivity during aging

We next examined the spatiotemporal dynamics of cross-compartment relocalization by grouping events according to the age interval in which each protein first switched compartments, yielding a directed “flow” map of inter-compartment redistribution across replicative aging (Figure S4A). Across the lifespan, most major compartments participated in bidirectional exchange—both gaining and losing proteins—yet each transition was dominated by a limited set of routes (Figure 4A, B, Table S3). Early aging was characterized primarily by nucleolus-to-nucleus relocalization, accompanied by prominent exchange among several other compartments. Between ages 3-5 to 6-8, nucleolus-to-nucleus flow remained strong and a mitochondria-to-cytosol wave emerged. From ages 6-8 to 9-11, the cytosol became a central hub, receiving proteins from multiple sources, including mitochondria, ER, and nucleus, while nucleolus-to-nucleus traffic persisted. In later transitions, the network narrowed to a smaller set of dominant flows, with continued nucleolar redistribution from ages 9-11 to 12-14 and increased influx from multiple compartments into the cytosol between ages 12-14 and 15-17. In the final transition (ages 15-17 to 18), relocalization was again dominated by nucleolus-to-nucleus redistribution, along with mitochondria and ER relocalization to cytosol. Together, these maps reveal recurring age-associated routes linking the nucleolus, nucleus, cytosol, and mitochondria, with nucleolar redistribution and multi-organelle-to-cytosol transitions emerging as dominant motifs.

**Figure 4.**
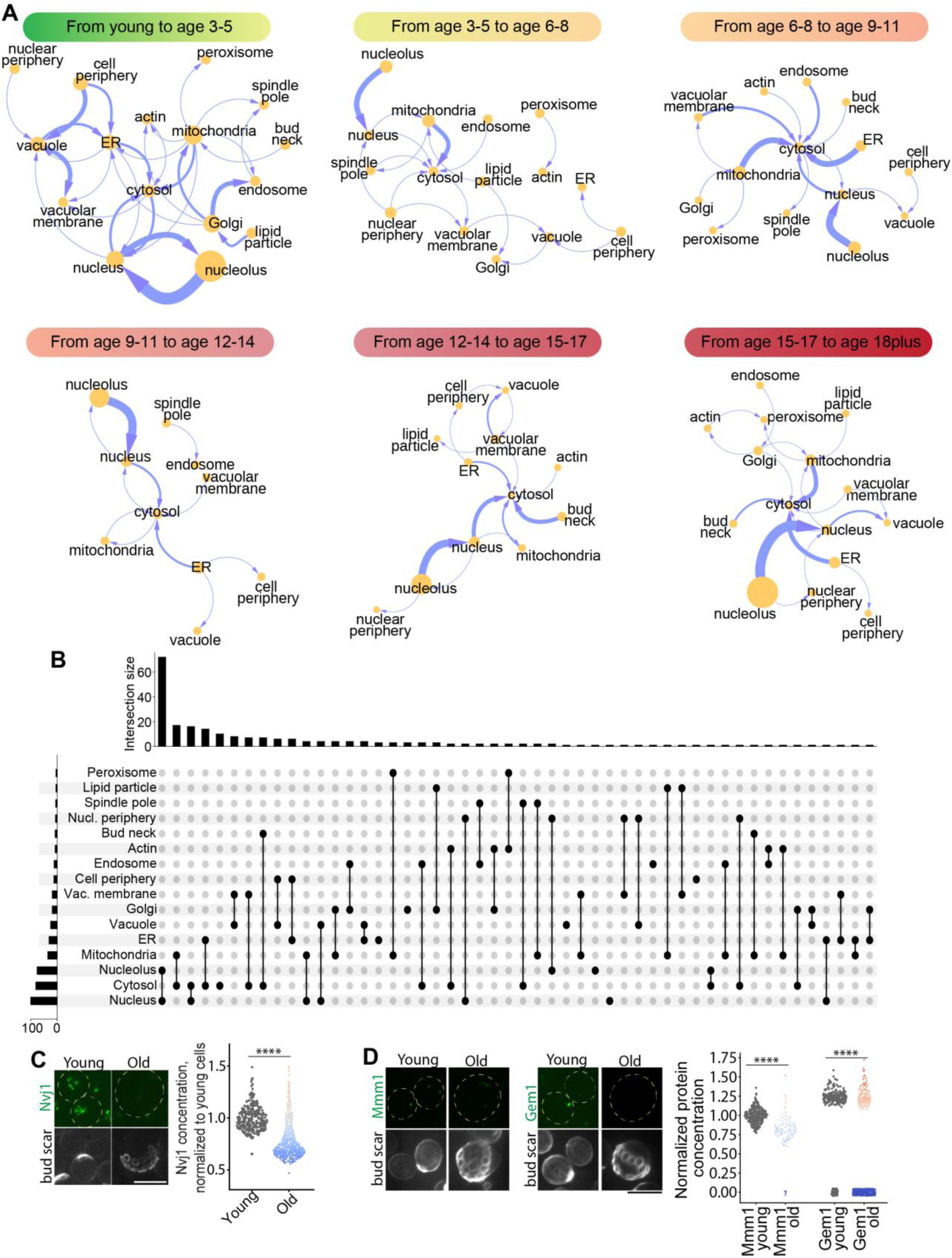
The landscape of inter-compartmental protein relocalization and compartment connectivity during aging. (**A**) Flow network of inter-compartmental protein relocalization during aging. Each node represents a protein localization class based on annotations in young cells, and node size reflects the number of proteins in each class. Arrow thickness indicates the number of proteins shifting from one compartment to another. **(B)** UpSet plot summarizing inter-compartmental protein relocalization events during aging in (A). Each set corresponds to a compartment in which proteins are detected in young cells, and intersections represent proteins that move between a defined pair of compartments (from their annotated location in young cells to a different compartment in aged cells). **(C**, **D)** Representative images and quantification of nucleus-vacuole contact (C) and mitochondria-ER contact (D) proteins during aging. Scale bar: 5μm. See Source Data table for number of cells quantified.

Because many of these relocalization events involve organelles connected by membrane contact sites, we next asked whether membrane contact-site components also change with age. As the catalog of contact-site proteins continues to expand, we focused on a small set of canonical interfaces—nucleus–vacuole, ER–mitochondria, and plasma membrane–cortical ER—as representative examples.

<u>Nucleus-vacuole interaction.</u> The nucleus-vacuole junction (NVJ) physically links the outer nuclear membrane to the vacuolar membrane and serves as a hub for sterol and neutral-lipid metabolism at ER–vacuole contacts^45,46^. NVJ formation depends on the interaction between Nvj1 (outer nuclear membrane) and Vac8 (vacuolar membrane)^47^. During aging, Vac8 remained expressed and properly localized to the vacuolar membrane, whereas Nvj1 was lost in more than half of cells early in aging, indicating that NVJs were impaired in a substantial fraction of cells (Figure 4C, S4B). Along with age-dependent mislocalization of ER lipid enzymes (e.g., Erg1, Hfd1, Ice2) discussed above, these NVJ defects are consistent with progressive disruption of a coordinated lipid network that normally channels sterols and lipids among these organelles.

<u>Mitochondria-ER interaction</u>. ER-mitochondria contacts support lipid exchange, calcium signaling, and mitochondrial dynamics^48^. In yeast, these contacts are mediated by the ER–mitochondria encounter structure (ERMES) complex (Mmm1, Mdm10, Mdm34, Mdm12) and regulated by the GTPase Gem1, which modulates contact-site distribution and mobility^48^. With age, some ERMES components showed changes in abundance and organization: Mmm1 puncta intensity declined markedly, and Gem1 puncta became undetectable, consistent with a progressive loss of stable ER–mitochondria tethering (Figure 4D). These changes suggest that aging compromises ER–mitochondria contact integrity, potentially disrupting lipid transport and metabolic coordination between these organelles.

<u>Plasma membrane-cortical ER interaction</u>. Plasma membrane (PM)–cortical ER contacts mediate communication between the cell surface and the ER and are established by Tcb1/2/3 and the membrane proteins Ist2 and Scs2^42^. These sites support lipid exchange, calcium signaling, and stress buffering at the cell periphery^42,49,50^. During aging, we observed a marked reduction in Tcb2/3 and Ist2 levels, accompanied by the formation of Scs2 foci, indicating progressive disorganization of PM–ER contact architecture (Figure S4C, S3H). These alterations imply declining PM–ER connectivity that may compromise lipid homeostasis, calcium signaling, and plasma membrane maintenance in aged cells.

Collectively, these results suggest that increased cross-compartment relocalization during aging reflects weakening of compartmental integrity and inter-organelle communication rather than enhanced coupling between compartments. Such decoupling—together with widespread protein mislocalization—may compromise coordinated organelle function during aging.

### Age-associated changes in organelle sizes

Beyond protein localization and inter-compartment communication, the relative sizes of major organelles—including the nucleus, vacuole, and mitochondria—set key functional ratios (e.g., transcriptional capacity, degradative/storage capacity, and ATP-producing volume) that shape coordinated organelle function within the cell’s crowded cytoplasm. We therefore quantified how organelle sizes change with age both in absolute terms and when normalized to cell size, and how their inter-organelle scaling relationships shift over the course of aging. In absolute terms, nearly all organelles increased in size, indicating that age-associated cell enlargement drives broad organelle expansion (Figure S5A). However, after normalizing to cell size, most organelles decreased in cell-normalized size, with vacuoles the only compartment that significantly increased relative to cell size (Figure 5A). Previous studies have shown that a reduced nucleus-to-cytoplasm ratio during cell size increase promotes cellular senescence in human cells^51,52^. Consistent with this, we observed a decline in cell-normalized nuclear size during replicative aging (Figure 5A), raising the possibility that sublinear scaling of the nucleus and other organelles relative to cell growth (i.e., organelle “dilution” relative to cell growth) is a conserved feature of cellular aging.

**Figure 5.**
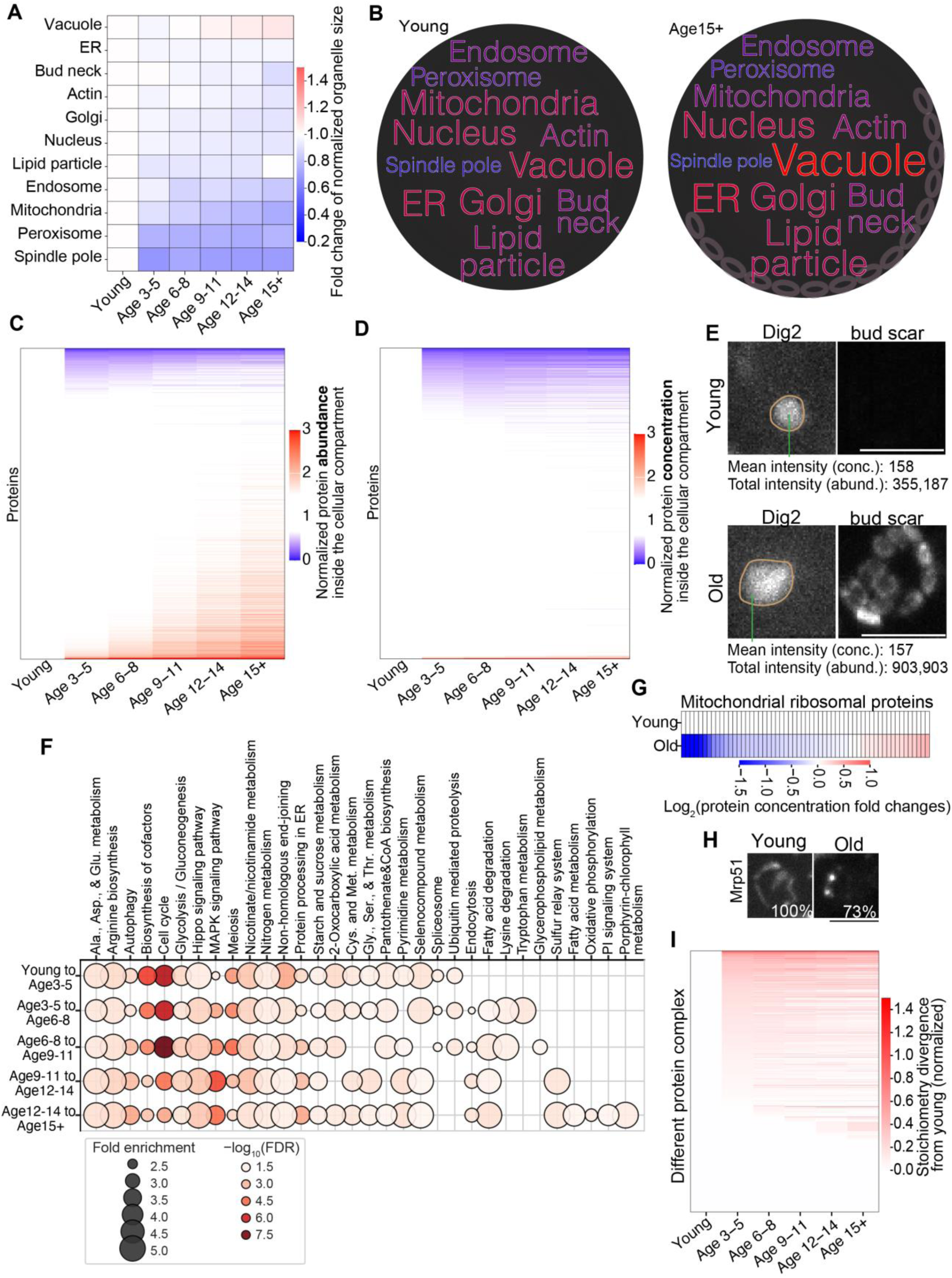
Age-associated changes in organelle size and protein concentration. (**A**) Age-associated changes in organelle size as a fraction of cell size. Organelle volumes were expressed as a fraction of cell volume at each age and plotted as fold change versus young. **(B)** Word cloud summarizing age-dependent changes in relative size among organelles. Organelle volumes were normalized to the total organelle volume (sum across organelles) to capture relative scaling behavior between organelles across age groups. **(C**, **D)** Proteome-wide quantification of protein abundance (C) and local concentration (D) within each compartment. Protein concentration was calculated by normalizing protein abundance to the measured size of the corresponding compartment, thereby distinguishing true concentration changes from apparent increases driven by organelle expansion or cell growth. For visualization, a 1.5× cutoff was used to highlight proteins exhibiting substantial age-associated change in abundance or concentration. **(E)** Representative images and quantification of Dig2 abundance and concentration in the nucleus during aging. Conc., concentration; abund., abundance. Scale bar: 5μm. **(F)** KEGG pathway enrichment for the proteins demonstrate significant concentration changes between different age groups. **(G**, **H)** Representative images and quantification of mitochondrial ribosome protein concentration changes during aging. Scale bar: 5μm. **(I)** Proteome-wide quantification of age-associated changes in protein complex stoichiometry. For each annotated complex, subunit concentrations were compared across age groups to assess whether the relative ratios among subunits remained constant or became imbalanced during aging. Complexes with coordinated changes across subunits indicate preserved stoichiometry, whereas disproportionate increases or decreases in specific subunits indicate stoichiometric remodeling and potential disruption of complex assembly or function.

Although most organelles decrease in cell-normalized size during aging, the timing of these changes differs across organelles: peroxisomes and the spindle pole body showed earlier reductions than mitochondria and endosomes, followed by other compartments (Figure 5A). Moreover, the extent of reduction was not uniform, leading to age-dependent shifts in inter-organelle proportions. For example, mitochondria and peroxisomes shrink much more than the nucleus, altering their scaling relationships over time (Figure 5B). Such loss of homeostasis in organelle size and scaling relationships may compromise coordinated organelle function and contribute to the metabolic remodeling observed during aging^15,53–55^.

### Global proteome abundance and local concentration changes during aging

The age-dependent changes in organelle size raised the question of whether protein amounts and concentrations within organelles scale proportionally. As a baseline, we first quantified whole-cell protein abundance across age, using whole-cell segmentation to measure total mNG signal per cell. By this metric, most proteins increased in abundance with age (2,717 increased versus 5 decreased with a 1.5-fold cutoff) (Figure S5B). Because aging alters both cell and organelle size—and because protein function depends on local concentration within compartments—it is important to distinguish changes in total cellular abundance from changes in compartmental abundance and concentration. Leveraging the subcellular resolution of our imaging dataset, we next quantified compartment-specific protein abundance and concentration dynamics in situ. To this end, we measured the mean mNG intensity within each segmented compartment as a proxy for local protein concentration using a local-thresholding approach ^12^.

Consistent with the whole-cell analysis, we observed widespread increases in the total organelle-localized protein amount (1,502 proteins increased and 303 decreased by >1.5-fold; Figure 5C, S5C, Table S4). The difference in the number of proteins showing an increase in abundance by whole-cell versus organelle-localized measurements likely reflects, in part, age-associated mislocalization, as exemplified above (e.g., Leu4). Surprisingly, within-compartment mean intensities changed more modestly and even in opposite directions, with 51 proteins increasing and 677 decreasing (Figure 5D, S5D, Table S4), indicating that some organellar proteins did not scale proportionally with organelle expansion, leading to reduced local concentrations (Figure S5D). Together, these patterns are consistent with organelle expansion accounting for much of the increase in organelle-localized protein amount (Figure 5E, S5A). These results highlight that measurements of total protein amount alone can overestimate functional changes when compartment size increases, whereas concentration better reflects the biochemical environment in which proteins operate. Functional enrichment of proteins with age-associated concentration changes highlighted pathways including amino-acid biosynthesis, cell cycle, glycolysis, and nicotinamide metabolism (Figure 5F, Table S4).

Although population-average concentrations were relatively stable for many proteins, aging increased cell-to-cell heterogeneity for a subset, with individual cells exhibiting divergent expression states (Figure S5E). For example, Nhp6b, a high-mobility group (HMG) chromatin protein, showed large increases in some aged cells and decreases in others, despite relatively modest changes in the population mean (Figure S5F). Similar behavior was observed for 541 proteins that exhibited significant concentration alterations in >30% of aged cells without a corresponding change in the population mean and were therefore not captured by analyses based on population-average local concentration (Figure S5G, Table S4). Notably, four proteins showed both increased and decreased concentration in substantial fractions of aged cells, consistent with divergent aging trajectories across individual cells (Figure S5H)^56^.

Importantly, age-associated concentration changes altered the stoichiometry of multi-protein complexes. For example, the mitoribosome (74 proteins in young cells) displayed heterogeneous subunit behavior: several components (e.g., Mrps8, Mrpl31, Mrpl32) decreased markedly, whereas many other mitoribosomal proteins were unchanged or increased (Figure 5G). For some increasing subunits (e.g., Mrp51), the local concentration rise coincided with increased aggregation, suggesting that a subset of “increases” may reflect punctate accumulation (Figure 5H). In addition to mitoribosomes, many other macromolecular complexes exhibited age-dependent shifts in subunit stoichiometry (Figure 5I, Table S4), suggesting widespread reorganization of proteome and protein interaction networks.

### Age-associated changes in protein associations

To test whether these stoichiometry shifts reflect broader remodeling of protein associations during aging, we used Protein Image-based Functional Annotation (PIFiA)^57^, an image-based approach to infer protein associations (including complexes and protein-protein interactions), and complemented it with limited proteolysis–mass spectrometry (LiP–MS), a biochemical readout of protein surface accessibility that is sensitive to changes in interaction interfaces.

PIFiA is a framework designed to infer protein associations from the feature profiles learned from single-cell imaging data by deep learning models^57^. To detect age-dependent changes in protein associations, we applied PIFiA to the high-dimensional feature profiles extracted from our 3D images by the ensemble localization model. Here, a **feature profile** refers to the learned multidimensional descriptors of 3D morphology and texture of the signal within each cell, and proteins with similar feature profiles often belong to the same macromolecular complexes or correspond to known interaction partners^57^. The ensemble model extracts a 2,560-dimensional feature embedding per cell that captures subtle morphology and texture information beyond compartment labels. We first benchmarked this approach by projecting feature profiles of different nucleolar complexes into a t-SNE map^57^. In this representation, proteins with similar feature profiles cluster together^57^. Indeed, subunits of the same complex form well-defined clusters that are separated from other complexes (Figure 6A, top), despite appearing similar by conventional localization patterns (Figure 6A, bottom). This benchmark confirms that, consistent with a previous report^57^, these subtle, high-dimensional feature profiles extracted by the model from 3D single-cell images are sufficient to discriminate distinct protein assemblies. We observed similar complex-dependent clustering for additional protein complexes across multiple compartments (Figure S6A–B), supporting the ability of image-derived feature profiles to resolve native assemblies from images^57^.

**Figure 6.**
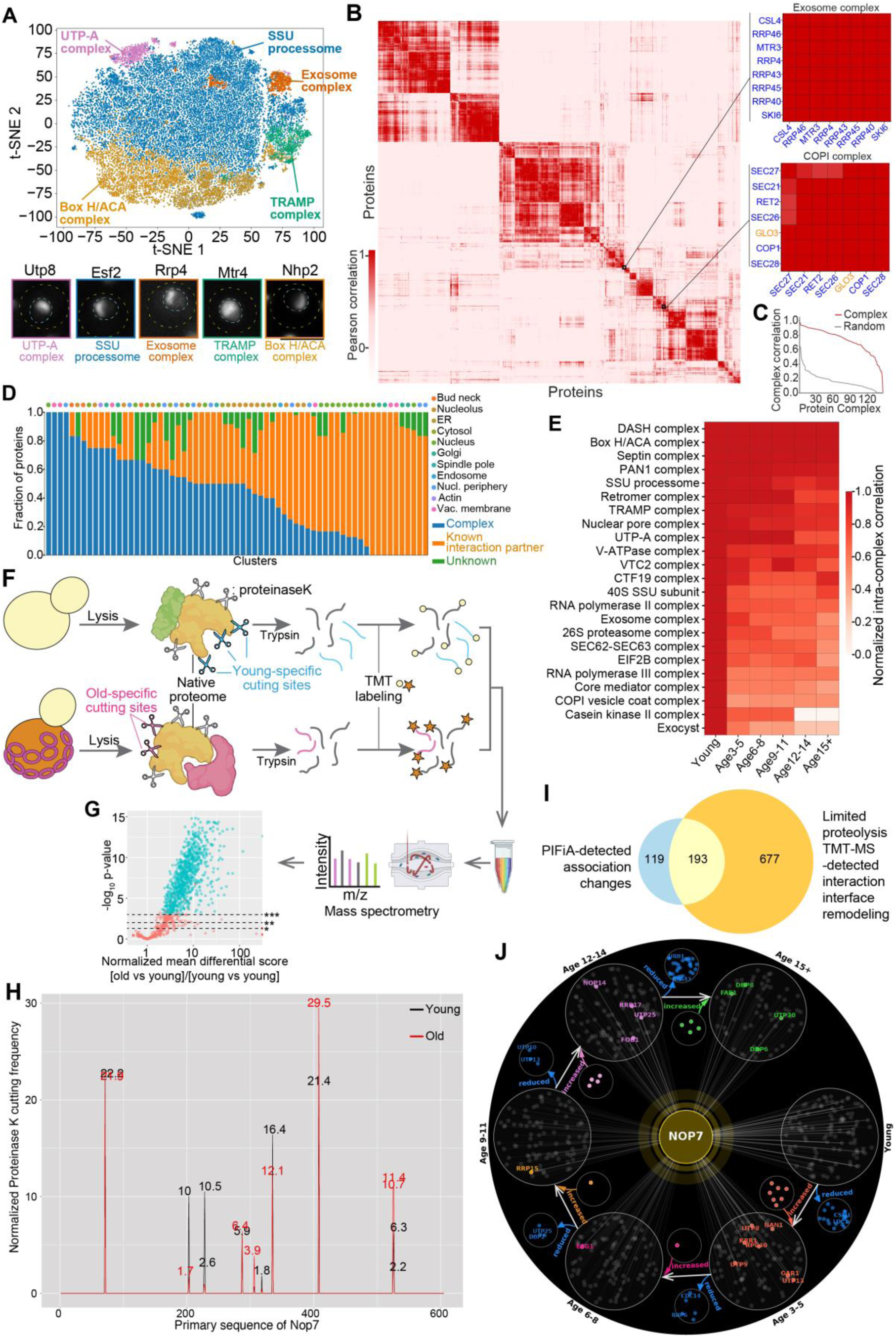
Age-associated changes in protein associations. (**A**) Single-cell tSNE plot based on PIFiA feature profiles and representative images of different nucleolar protein complexes. Each dot is a single cell; cells from different subunit proteins of the same complex are colored the same. Note that these different nucleolar complexes exhibit similar subcellular localization by eye (bottom), yet display visually imperceptible differences that are captured by PIFiA feature profiles and used to distinguish the clusters (top). Scale bar: 5μm. **(B**, **C)** Proteome-wide pairwise correlations of PIFiA feature profiles (B) and performance of PIFiA feature profile-based clustering for protein complexes and associations (C) from young cells. Insets highlight correlations among members of two example complexes (the exosome and the COPI complex). In these zoomed views, blue gene names indicate annotated core subunits of the complex, whereas orange gene names indicate proteins known to interact with the complex. (C) compares the mean within-complex correlation (complex; red line) to the mean correlation of randomly sampled protein sets matched for complex size (random; gray line). (**D**) Fraction of proteins in each PIFiA cluster of young cells that are reported as members of a protein complex (blue) or have documented protein–protein interactions with other proteins in the same cluster (orange). Proteins with no known interactions with other proteins in their cluster are shown in green. The predominant subcellular localization for proteins in each cluster is indicated above. (**E**) Age-associated changes in PIFiA feature-profile similarity among known subunits of protein complexes identified from the clusters in (D). (**F**) Workflow of limited proteolysis-TMT mass spectrometry to assess the age-associated changes of protein associations via surface accessibility profiling. (**G**) Volcano plot of proteome-wide, age-associated changes in differential score (ANOVA). Age-associated changes were normalized to the differential-score variation observed between biological replicates of young cells. Proteins showing at least 2-fold age-associated changes in differential score are highlighted in cyan. *: p<0.05; **: p<0.01; ***: p<0.001. (**H**) Example protein Nop7 showing age-associated changes in proteinase K accessibility across its primary sequence. Numbers above peaks indicate the peak height (normalized cutting frequency) for each condition, enabling comparison when young and old peaks overlap at the same sequence position. (**I**) Venn diagram showing the overlap between protein association changes detected by PIFiA feature profiles and by limited-proteolysis TMT mass spectrometry. (**J**) PIFiA feature profile-inferred, age-associated remodeling of Nop7’s previously reported interaction network. Each surrounding sector corresponds to a defined age bin. Within each age bin, each node connected to Nop7 represents a previously reported interaction partner whose PIFiA feature profile shows significant similarity to Nop7. Comparisons across age bins highlight partner-specific gains and losses of co-clustering/feature-profile similarity with Nop7, indicating dynamic remodeling of Nop7’s interaction network. Proteins with increased versus decreased similarity to Nop7 are highlighted separately, revealing age-dependent strengthening or weakening of specific associations.

We next evaluated the proteome-wide ability of PIFiA to recover previously reported protein complexes and associations across compartments (Figure S6C)^57^. Clustering based on pairwise feature-profile similarity (hereafter “PIFiA clusters”) revealed high correlation between proteins that form known protein complexes (Figure 6B-C, S6C). As expected, in addition to resolving established protein complexes, PIFiA analysis also recovered additional proteins that are not annotated as core complex subunits but cluster tightly with known complexes (Figure 6B, 6D). Consistent with prior work^57^, these co-clustering proteins often correspond to well-known, functionally associated interaction partners of the complex. For example, Glo3, which binds and regulates COPI, co-clustered with COPI proteins (Figure 6B). Similar patterns were observed across multiple compartments, indicating that this feature profile-based clustering can recover both complex subunits and closely associated interaction partners (Figure 6D)^57^. In the following analyses, we restricted PIFiA feature profile comparisons to literature-curated protein complexes/associations and used changes in PIFiA co-clustering/feature-profile similarity among known partners as a readout of association changes.

We then compared these high-fidelity clusters between young and old cells to examine how previously reported protein complexes and known interaction partners change with age. This analysis revealed extensive remodeling of complexes and their interaction networks during aging, evident as a broad weakening of feature-profile co-clustering among complex subunits and interaction partners in aged cells (Figure 6E). In some cases, this was also accompanied by stoichiometric imbalances among complex subunits and shifts in subcellular distribution of specific proteins relative to their partners. For example, proteasome subunits showed unequal reduction during aging; at the same time, components such as Rpn5 and Rpn9 relocalize to the cytosol, whereas most other subunits remain predominantly nuclear (Figure S6D).

To complement PIFiA-based analysis of age-dependent changes in protein associations, we applied limited proteolysis coupled to tandem mass tag (TMT)–based quantitative mass spectrometry (LiP–MS) to profile proteome-wide alterations in protein surface accessibility^58,59^. Limited proteolysis with proteinase K (PK) under native conditions preferentially cleaves solvent-exposed sites, providing a sensitive readout of changes in surface protection that can arise from altered protein associations (Figure 6F)^58,59^. We therefore compared young and aged cells by LiP–MS, analyzing soluble lysates to focus on non-aggregated native proteins, and quantified PK-generated peptides by TMT-based mass spectrometry (Figure 6F). We summarized these data as differential PK accessibility scores for each protein between young and aged samples (Figure S6E). This workflow was highly reproducible, with only minor differences in surface accessibility among biological replicates (Figures S6F–G). In contrast, we observed robust shifts in surface accessibility between young and aged samples, reflected by large changes in differential accessibility scores (Figures S6G–H).

Using this pipeline, we identified over 800 proteins exhibiting significant changes in surface accessibility, consistent with widespread alterations in protein interaction interfaces (Figure 6G, Table S5). For example, Nop7 showed minimal variation in surface accessibility across young cell replicates but displayed pronounced deviations in aged cells (Figure 6H, S6I), whereas Num1 exhibited little to no difference between age groups (Figure S6J). Given incomplete proteome coverage of mass spectrometry, this set likely underestimates the full extent of age-dependent remodeling of interaction interfaces. Importantly, these age-induced shifts in surface accessibility do not simply reflect amplification of the baseline heterogeneity already present among young cells (Figure S6H). Moreover, the magnitude of surface-accessibility changes did not correlate with protein melting temperatures or predicted disorder scores, indicating that these alterations are unlikely to simply reflect nonspecific misfolding of intrinsically labile proteins (Figure S6K-L). Proteins showing age-dependent changes in surface accessibility were enriched in key biological processes, including amino acid biosynthesis, nuclear transport, ribosome biogenesis, and translational regulation (Figure S6M). Collectively, these data support widespread age-associated remodeling of protein interaction interfaces.

Integrating LiP–MS with our PIFiA-based inference of interaction-network changes, we found that ∼62% of protein pairs showing age-dependent shifts in PIFiA feature-profile similarity also exhibited significant changes in surface accessibility, supporting the robustness of these inferred association changes (Figure 6I, Table S5). For example, focusing our PIFiA comparisons on literature-curated complexes and associations, we found that Nop7 exhibited partner-specific gains and losses of co-clustering across ages, indicating age-dependent strengthening of some previously reported interactions and weakening of others (Figure 6J, Table S5). Consistent with this, Nop7 exhibited region-specific increases and decreases in proteinase K accessibility in LiP–MS (Figure 6H), supporting age-associated remodeling of its interaction interfaces. Similar patterns are observed for Rps6a (Figures S6N–P). Together, these results indicate that aging remodels protein complex stoichiometry and interaction networks for some proteins, likely contributing to rewired signaling and altered cellular functions in aged cells.

### The protein structural features underlying the age-associated change in the proteome

We next sought to understand the mechanisms underlying the observation that some proteins undergo pronounced age-associated remodeling whereas others remain comparatively stable. We first quantified the deviation of each protein’s localization scores at a given age relative to its baseline profile in young cells. The top 30% of proteins showing the strongest age-associated deviations were defined as the “unstable” group, while the bottom 20% with minimal or no detectable changes were designated as the “stable” group for downstream comparisons (Figure S7A). Additionally, proteins exhibiting more than 2-fold changes in concentration were included in the unstable group. We compared a range of basic physicochemical features derived from primary sequences—including protein length, hydrophobicity, fraction of disordered residues, and number of domains—between the unstable and stable groups. However, none of these classical biochemical features showed a clear correlation with a protein’s susceptibility to aging-associated changes (Figure S7B), indicating that primary-sequence features alone are insufficient to explain age-associated protein remodeling.

Next, we turned to protein three-dimensional structural properties as potential determinants of age susceptibility. To uncover deeper determinants, we leveraged a comprehensive set of 41 structural and biophysical features derived from the 3D structures of each protein predicted by AlphaFold (Figure S7C, Table S6). Using these features, we trained a logistic regression model on 690 unstable and 614 stable proteins (Figure 7A). Receiver operating characteristic (ROC) curve analysis indicated that this initial model captured key features underlying age-associated protein changes, achieving an area under the curve (AUC) of 0.86 (Figure S7D). Examination of feature contributions allowed us to remove redundant features and identified 28 key structural parameters that preserve high discriminative performance (Figure S7C-D). Importantly, random train/test splits of aging-stable and aging-unstable proteins yielded consistently high performance on held-out data, indicating that the structural signatures underlying age susceptibility are robust and generalizable across proteins (Figure 7B, S7E).

**Figure 7.**
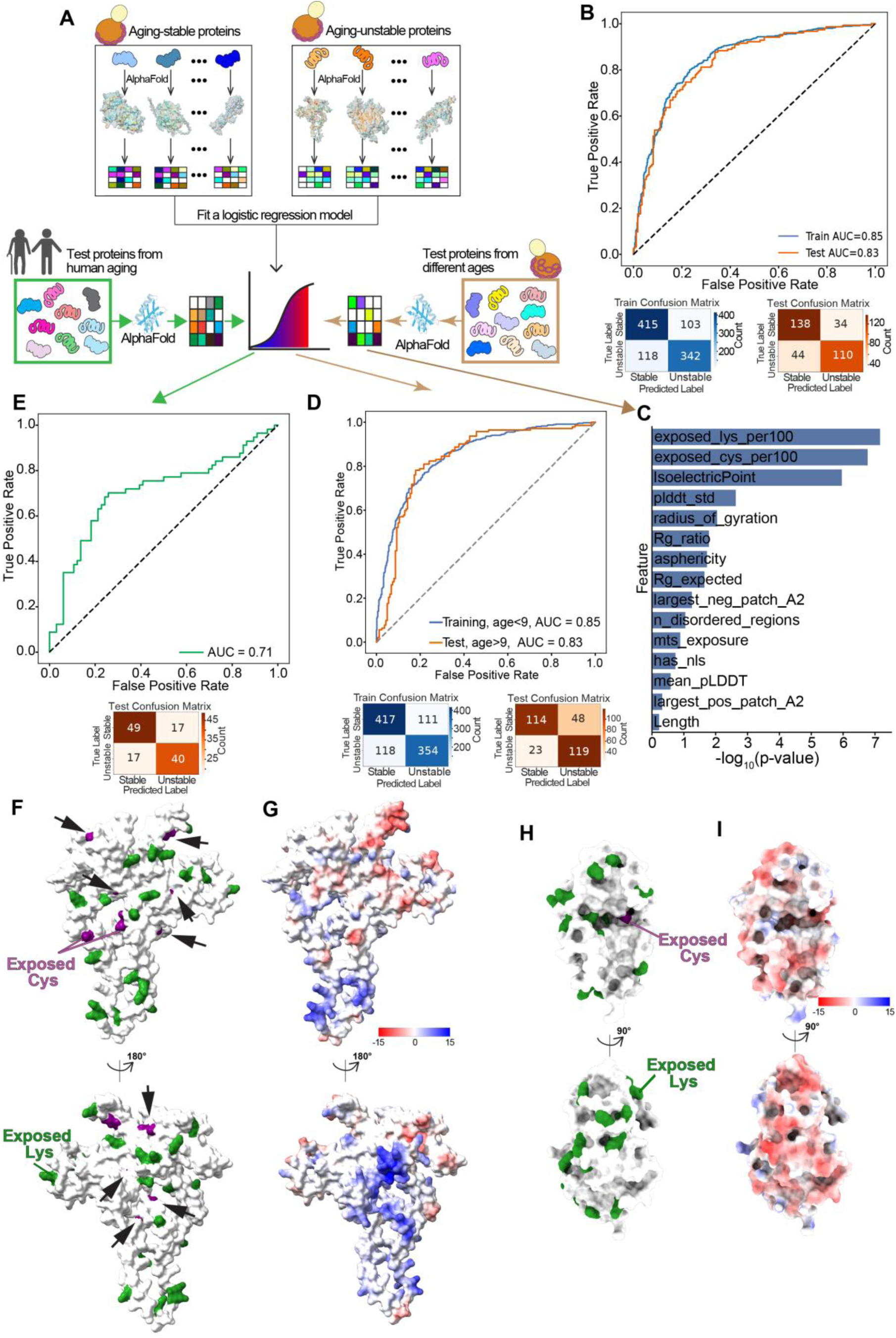
Intrinsic structural determinants of protein fate during aging. (**A**) Schematic of the logistic regression framework used to distinguish aging-stable and aging-unstable proteins. Proteins were classified as stable or unstable based on age-associated changes in localization scores or protein concentration, and a panel of physicochemical/structural features was extracted from their AlphaFold-predicted 3D models. These features were used to train a logistic regression model to discriminate aging-stable from aging-unstable proteins. To test whether shared features underlie proteome remodeling across the lifespan, stable/unstable protein sets from different yeast aging stages in this study were used as training and testing data (bottom right of (A)). In addition, the model trained on the yeast proteome was applied to a published human dataset to evaluate cross-species generalization of the identified physicochemical/structural determinants (bottom left of (A)). **(B)** ROC curve analysis evaluating the ability of structural and physicochemical features to predict experimentally defined aging-stable versus aging-unstable proteins using a logistic regression model trained on proteins classified as stable/unstable across all age stages. These proteins were randomly split into a training set and a held-out testing set. The corresponding confusion matrices for the training and held-out testing datasets are shown below. **(C**, **D)** (C) Top structural/physicochemical features most strongly associated with aging instability in the held-out test set of proteins from cells aged ≥9 generations. (D) ROC curve analysis assessing the ability of these structural/physicochemical features to predict aging-stable versus aging-unstable proteins in age ≥9 cells using a logistic regression model trained on earlier age groups. The corresponding confusion matrices for the training and held-out testing datasets are shown in (D). **(E)** ROC curve analysis evaluating the ability of structural and physicochemical features of human proteins to predict their aging-instability using a logistic regression model trained on the yeast aging-stable/unstable proteome. The corresponding confusion matrix for human testing dataset is shown below. **(F**-**I)** Exposed cysteine (purple)/lysine (green) residues (F, H) and surface charges (G, I) mapped onto AlphaFold models of Ice2 (F, G) and Rki1 (H, I). Black arrows indicate exposed Cys residues; Rki1 contains only a single exposed Cys residue. Structural models were obtained from AlphaFold (AF-P40499-F1-model_v6 and AF-Q12189-F1-model_v6).

Encouraged by these results, we next tested whether a model trained on proteins showing age-associated changes early in aging could predict those occurring later. We divided proteins with age-associated changes into two temporal classes: early-aging proteins (changes within the first 8 divisions) and late-aging proteins (changes after 9 cell divisions). This yielded 528 stable and 472 unstable proteins during the first eight cell divisions, which we used to retrain a logistic regression model with the same 28 features as above; the resulting classifier accurately captured early-aging behavior with an AUC of 0.85 (Figure 7C–D). We then applied this model to predict the fates of 162 stable and 142 unstable proteins that show age-associated changes after nine cell divisions using their structural features, achieving a robust AUC of 0.83 (Figure 7D, Table S6). These results indicate that similar structural principles govern protein susceptibility to aging across different stages of the lifespan, motivating us to test this model in other species, as structural features are highly conserved.

To assess the generalizability of our yeast-trained model, we evaluated it on a human proteomic dataset of age-associated changes derived from large-scale mass spectrometry. Because the human proteome exhibits substantial tissue-specific variability, we focused on UUPAs (ubiquitously upregulated proteins with aging) and UDPAs (ubiquitously downregulated proteins with aging) identified in a recent study^60^. These proteins, which are consistently increased or decreased across multiple tissues in older individuals, were classified as the unstable group, whereas broadly expressed proteins with minimal variability and no age-associated change were designated as the stable group. We extracted the same 28 AlphaFold-derived structural features for these human proteins and applied the yeast-trained model to classify them based on their structural features. The model robustly reproduced the experimental classification, achieving an AUC of 0.71 (Figure 7E). Notably, it correctly labeled 27 of 28 UUPAs as unstable (Table S6), indicating that these proteins share similar structural signatures that predispose yeast proteins to age-related instability. This slightly reduced accuracy, especially for UDPAs, may reflect methodological differences, as mass spectrometry–based abundance is inferred from detected peptides; age-dependent post-translational modifications (PTMs) and isoform shifts (including alternative splicing) can alter or mask peptide sequences, reducing peptide recovery/identification and thereby biasing abundance estimates downward even when total protein is stable^61–63^. Together, these findings indicate that intrinsic structural features are key determinants of protein susceptibility to aging across evolution and demonstrate that our imaging-derived yeast proteome dataset captures fundamental principles of proteome aging that generalize across species and measurement modalities.

The most significant features underlying our model’s predictions of age-associated protein changes included the number of surface-exposed cysteine and lysine residues, isoelectric point, pLDDT scores, and radius of gyration (Figure 7C). Surface-exposed cysteines and lysines emerged as the strongest predictors, consistent with the high chemical reactivity of their thiol and ε-amino side chains and their propensity for enzymatic and non-enzymatic PTMs^64–67^. With aging, shifts in redox homeostasis are likely to render cysteine-rich proteins particularly vulnerable, while PTMs on cysteine and lysine can alter protein activity and stability and remodel interaction interfaces and complex assembly. Consistent with this view, isoelectric point further tunes protein surface properties and interaction propensity. In addition to surface properties, features such as pLDDT and radius of gyration likely reflect overall protein stability and compactness: lower pLDDT values and a larger radius of gyration are consistent with less ordered and/or less compact proteins that may be more susceptible to age-dependent destabilization. These structural signatures are exemplified by the age-associated unstable protein Ice2, which is enriched in exposed cysteines and lysines, has a high isoelectric point due to enriched surface charges, exhibits lower pLDDT scores, and undergoes an age-dependent shift in localization (Figure 7F–G, S3H, S7F). In contrast, the aging-stable protein Rki1 has only one surface-exposed cysteine, a low isoelectric point, high pLDDT scores, and much higher compactness (Figure 7H–I, S7F-G).

## Discussion

We generated a comprehensive, proteome-wide dataset that captures both protein concentration and spatial organization changes throughout the replicative aging of budding yeast. This dataset provides single-cell, single-age resolution of proteome dynamics, encompassing changes in protein concentration, localization, interaction, and aggregation. The spatiotemporal imaging data are further integrated with LiP-MS in young and aged cells, enabling detection of aging-associated remodeling of interaction networks. The breadth and resolution of this dataset provide a unified foundation and reference for systematic hypothesis generation and mechanistic discovery in the biology of aging. Moreover, this imaging and computational framework can be readily extended to spatiotemporal analyses of the proteome under diverse longevity paradigms and environmental exposomes, offering a scalable approach to dissect how cellular and environmental factors shape the trajectory of aging.

The observation that intrinsic structural features of proteins can predict their vulnerability during aging is both surprising and conceptually significant. Classical aging theories, including the disposable-soma hypothesis, emphasize the stochastic accumulation of molecular damage that broadly impairs macromolecular function. While such models help explain the heterogeneity of aging phenotypes among cells and individuals, they do not fully account for the reproducible molecular hallmarks and biomarkers of aging observed at the population level. Our findings introduce an additional structure-linked layer: intrinsic protein structural features predispose specific proteins to aging-associated remodeling. Surface properties such as exposed cysteine or lysine residues, local charges, and structural flexibility can render certain proteins particularly sensitive to redox and other physicochemical alterations in aging cells. Consequently, each protein may carry a defined intrinsic susceptibility to the aging environment, leading to dysfunction at characteristic rates and emerging as biomarkers.

This structure-encoded vulnerability provides a unifying framework linking stochastic damage at the level of individual cells to reproducible population-level patterns, in which particular proteins and complexes are consistently affected during aging. It also resonates with the antagonistic pleiotropy theory, wherein structural features of some proteins that confer adaptive advantages early in life—such as catalytic reactivity, protein-protein interactions, or dynamic flexibility—become liabilities later, as these same features increase susceptibility to oxidative and metabolic stress. Thus, the architecture of the proteome encodes both functional potential and fragility, defining an intrinsic aging trajectory shaped by evolutionary trade-offs between performance and stability. The conservation of these structural determinants across species further supports the notion that the proteome’s design constrains how aging unfolds across evolution. In this view, aging preferentially perturbs network nodes defined by structural vulnerability, providing a mechanistic bridge between molecular biophysics and systems-level manifestations of cellular aging.

## Acknowledgments

We thank Dr. Maya Schuldiner and Dr. Michael Knop for providing SWAT libraries. We also thank the input and comments from lab members and colleagues, especially Catherine Chang. This work was supported by DP5OD024598, R21AG077556, Impetus grant, Larry L. Hillblom Foundation 2023-A-007-SUP, and R01AG075201 to C. Zhou and Hevolution Foundation to the Buck Institute for Research on Aging (HF-PART-23-1422047).

## Author contributions

C.Z. conceived and supervised the project. S.Y., L.L., and C.Z. designed the research. S.Y. and L.L. performed proteome-wide imaging data collection. L.L. and L.V. developed the machine-learning pipelines. L.V. analyzed protein localization, and L.L. analyzed proteome-wide protein abundance and concentration. L.V., L.L., and C.Y. performed data analysis and generated figures. Q.L. prepared LiP–MS samples, which were analyzed by Z.W., Y.Z., and L.F.. J.Z. and F.Z. generated plasmids and strains for localization validation. K.S. provided support for the MS. C.Z. drafted the manuscript with input from all authors.

## Yeast strains and culture condition

All *S. cerevisiae* strains, plasmids, primers, antibodies, and chemical reagents used in this study are listed in Key Resource Table. Yeast strains used in this study are based on the BY4741 strain background. Genetic modifications were performed with PCR mediated homologous recombination (Longtine et al., 1998)^81^ and genotyped with PCR to confirm correct modification and lack of aneuploidy for the chromosome that gene of interest is located. The mNG-tagged strains were from the seamless tagging collection (Meurer et al., 2018)^16^. All plasmids were constructed based on Gibson assembly with Gibson Assembly Master Mix (New England Biolabs, E2611S). Expression of proteins from integration plasmids was done by integrating the linearized plasmid into TRP1 locus.

Yeast cells were grown at room temperature in YPD (10 g/L yeast extract from BD, 20 g/L peptone from BD, and 20 g/L glucose), and refreshed for additional 2-3 hrs before imaging. The OD_600_ of cell culture was maintained between 0.4–1.0 overnight and throughout the experiments to avoid metabolic shifts caused by glucose exhaustion. All mediums used in the study were prepared by autoclaving the yeast extract and peptone for 20 min before adding the filtered carbon source as indicated.

## Confocal microscopy

Cells were imaged in 384-well glass-bottom plates (Cellvis, P384-1.5H-N, USA). Images were acquired on a Nikon CSU-W1 SoRa spinning-disk confocal microscope equipped with a 60×/1.27 NA Plan-Apochromat water-immersion objective (Plan Apo IR 60× WI DIC N2). Z-stack acquisition was controlled by a piezoelectric Z stage to ensure accurate and reproducible step spacing. Excitation was provided by 405, 488, and 561 nm laser lines to image calcofluor white, mNeonGreen (mNG), and mScarlet3, respectively; wheat germ agglutinin (WGA) was imaged using 561 nm excitation as indicated. Emission light was collected through bandpass filters and detected on an ORCA-Fusion BT sCMOS camera using the following channels: 455/50 nm (calcofluor white), 520/40 nm (mNG), 605/52 nm (mScarlet3), and 620/60 nm (WGA). For selected fields of view, images were acquired in SoRa mode to increase lateral resolution. Unless otherwise stated, all conditions within an experiment were imaged using identical acquisition settings (laser power, exposure time, Z-step size, and detector parameters) to enable quantitative comparisons. Raw image stacks were processed and analyzed using Fiji/ImageJ (NIH, Bethesda, MD).

## Proteome-wide old cell enrichment and imaging

<u>Automated old cell enrichment</u>: Automated old cell enrichment was performed using a biotin-streptavidin-based labeling strategy combined with automated Tecan Fluent 480 liquid-handling robot. Approximately OD₆₀₀ = 1.5 of mid-log phase yeast cells were transferred to 96-deep-well plates and washed three times with PBS buffer (pH 8.0). Cells were resuspended in PBS (pH 8.0) and incubated with 0.35 mg NHS-dPEG®12-biotin (Sigma, QBD10198) at room temperature for 30 min with intermittent mixing. Excess biotin was removed by four washes with PBS buffer (pH 7.2). Biotinylated cells were then mixed with 15 µL of BioMag streptavidin beads per strain (Polysciences, 84660-5) and incubated at room temperature with intermittent mixing for 30 min to allow bead binding. Bead-bound cells were collected using a 96-well plate magnet (Alpaqua, A000400), followed by replacement with fresh YPD medium. Replicative aging was carried out for 23 hours on the deck of Tecan robot, with medium replacement every 2 hours by transferring to 96-well magnet plate, while maintaining gentle resuspension to prevent bead aggregation and cell sedimentation by blowing air bubbles into the culture medium every 10 minutes. Before imaging, cells were stained with WGA594 (CF®594 Wheat germ agglutinin (WGA) conjugates) at 1 µg/mL for 1 hour to label bud scars, followed by Calcofluor White staining to outline the cell wall. For Calcofluor White staining, 0.7 µL of Calcofluor White stock solution (1 mg/mL; Sigma-Aldrich, 18909-100ML-F) was added to 1 mL of cell culture and incubated for 10 minutes. These cells were washed three times using magnetic plate to remove dyes. Except for cell collection before magnetic bead tagging, which was performed by low-speed centrifugation (1,600 xg), all washing, labeling, staining, and culturing steps were performed using an automated Tecan Fluent 480 liquid-handling system.

<u>Automated imaging</u>: To reduce the time cells spent waiting on the microscope stage, the 96 strains were processed in two batches: enriched old cells from 48 strains were washed and stained for imaging in each batch. Live enriched aged cells were then transferred to 384-well glass-bottom plates and imaged automatically on a Nikon CSU-W1 SoRa spinning-disk confocal microscope equipped with a 60× water-immersion objective (NA 1.27), an ORCA-Fusion BT sCMOS camera, and a piezo Z-stage for rapid and precise axial positioning. The entire acquisition workflow was automated using the NIS-Elements JOBS module. For each well, autofocus was performed on the Calcofluor White channel using a software-based routine. Z-stacks were acquired using hardware triggering to maximize acquisition speed and minimize mechanical vibration during high-throughput live-cell imaging.

For each well, images were collected from 25 fields of view (FOVs) in sequential channel order (mNG, WGA, then Calcofluor White). For the mNG and Calcofluor White channels, 9 focal planes were acquired with a 1.0 µm Z-step size; for the WGA channel, 21 focal planes were acquired with a 0.5 µm Z-step size. Standard (non-SoRa) acquisition settings were 45% laser power with 70 ms exposure (405 nm), 35% laser power with 300 ms exposure (488 nm), and 100% laser power with 100 ms exposure (561 nm). In total, about 47.5TB data was collected for the entire proteome.

## FM4-64 staining

For vacuole visualization, a 4 mM FM4-64FX stock was prepared by dissolving 100 μg of FM4-64FX (Thermo Fisher Scientific, F34653) in 32 μL DMSO. Cells were labeled at a 1:1,000 dilution in 1 mL YPD for 1 hour at 30 °C in the dark. For young cells, stained cultures were diluted into 10 mL fresh YPD and incubated for an additional 5 hours at 30 °C prior to imaging. For aged cells, the staining protocol was adapted to the aging workflow: after 20 hours of automated culture in the Tecan robot, cells were temporarily removed, stained in 1 mL YPD for 1 hour at 30 °C in the dark, then returned to deep-well plates and cultured for an additional 3 hours in the Tecan robot before imaging.

## Deep learning models and proteome-wide image analysis

<u>Cell segmentation.</u> Cells were segmented using Cellpose (v2.2.3) ^82^based on the Calcofluor White (CW) signal. The CW z-stacks were first maximum-intensity projected, and individual cells were segmented with the Cellpose ‘cyto’ model using a fixed diameter of 80 pixels. The resulting mask was then used to crop the corresponding GFP and WGA image stacks into 64 × 64 pixel single-cell image volumes for downstream analysis.

<u>Bud scar detection and counting with **DeepAge** model.</u> To determine total bud scar counts, WGA image stacks were partitioned into upper and lower hemispheres using a Hough circle–based equator detection pipeline. Each z-slice was processed via Canny edge detection (σ = 2; low threshold = 1; high threshold = 50) to generate binary edge maps. A circular Hough transform (radii 20–30 pixels) was then applied, retaining the top two circle peaks. The cell equator was designated as the z-slice with the highest equator score, defined as:

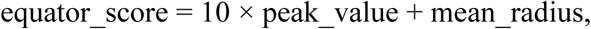

where peak_value is the primary accumulator value and mean_radius is the average radius of the two retained circles. Slices above and below the equator were maximum-intensity projected to generate discrete hemispherical images for independent analysis. Bud scars were detected in these projections using DeepAge, a deep learning-based object detection model built on the YOLOv8 framework. The model was fine-tuned for a single detection class using a manually annotated dataset of 441 training images. Training was performed for 100 epochs with a batch size of 16 using the AdamW optimizer, a weight decay of 5×10^−4^, and automatic mixed precision. Although native images were 64 × 64 pixels, they were rescaled to 640 × 640 during training and inference. To ensure robust performance, we utilized an early stopping patience of 50 epochs. Model was validated on 132 images (337 annotated instances). The final bud scar number for each cell was reported as the sum of detections from both the upper and lower hemisphere projections.

## Ensemble model

<u>Data Preprocessing:</u> A high-confidence ground-truth dataset comprising 37,816 manually annotated single yeast cells was constructed for protein subcellular localization. The dataset was partitioned using stratified random sampling to preserve class balance across splits: 81% training (29,820 cells), 9% validation (3,314 cells), and 10% held-out testing (3,682 cells). The test set was excluded from all stages of model development, including model selection and hyperparameter tuning, and was used exclusively for final performance evaluation.

Two complementary input representations were derived from the dataset to model intracellular organization at different spatial resolutions. The analysis pipeline comprised two parallel model streams, operating on 2D projection-based inputs and 3D volumetric inputs, respectively. The two model streams were trained independently and integrated at the decision level.

<u>2D classification model via transfer learning from DeepLoc</u>: For the 2D classification model, we followed the input conventions of the original DeepLoc model. Eight-slice z-stacks were collapsed using maximum-intensity projection (MIP) to generate 64 × 64 pixel 2D images. Pixel intensities were then normalized to the range [0, 1] by saturating the top and bottom 0.1 percentiles of the intensity distribution to ensure compatibility with the pre-trained DeepLoc weights.

The convolutional architecture is based on the DeepLoc model originally introduced by a previous work^18^ and subsequently adapted and re-trained for yeast proteome-scale localization analysis using transfer learning in our prior work ^59^. We adopted this validated implementation as the starting point for the present study.

The network consists of 11 layers organized into eight convolutional blocks followed by three fully connected layers, and is designed to process 64 × 64 pixel inputs. The convolutional feature-extraction layers (layers 1–10) were initialized with weights from the previously trained model. Inputs consisted of two channels (mNG maximum-intensity projection and a corresponding cell mask), consistent with the DeepLoc configuration. The network was fine-tuned using the combined manually annotated training and validation set (*n* = 33,134 cells), while an independent test set (*n* = 3,682 cells) was held out and used exclusively for final performance evaluation.

3D Volumetric Classification model based on 3D ResNet

*<u>Architecture of the model</u>*: To capture the full spatial context of intracellular organization, we developed a 3D volumetric classification model centered on individual cell crops. Each single-cell ROI was standardized into a volumetric tensor of 8 x 64 x 64 voxels (Depth x Height x Width). This specific configuration was chosen to preserve the native axial resolution of the microscope, utilizing all 8 collected optical slices without interpolative loss. Prior to network entry, voxel intensities underwent Z-score standardization (zero mean, unit variance) to mitigate batch-to-batch intensity variance and ensure stable gradient convergence by preventing internal covariate shift during the volumetric training phase.

The architectural backbone was a 3D Convolutional Neural Network (CNN) adapted from the ResNet-50 framework^19^. The network begins with a 7 × 7 × 7 convolutional stem with stride 1 followed by 3D max pooling, a design that preserves spatial resolution in volumetric inputs. The backbone comprises four residual stages containing 3, 4, 6, and 3 bottleneck blocks, respectively. Each bottleneck block consists of sequential 1 × 1 × 1, 3 × 3 × 3, and 1 × 1 × 1 convolutions with identity-based residual skip connections, enabling effective gradient propagation in deep networks. All convolutional layers are followed by batch normalization and ReLU activation, consistent with the original ResNet design. The network terminates in a global adaptive average pooling layer that aggregates spatial information into a 2048-dimensional feature vector, which is passed to a fully connected layer to predict 17 subcellular localization classes (16 organelle classes plus a “None” class corresponding primarily to unexpressed cells).

*<u>Training and optimization strategy</u>*: To mitigate overfitting and improve generalization to unseen morphologies, we applied on-the-fly 3D data augmentation during the training phase. Transformations included random spatial flips along all three axes (p=0.5), random 90-degree rotations in the XY plane, and random volumetric zooming (scaling factors 0.9x to 1.1x). These augmentations forced the network to learn features invariant to orientation and minor scale variations rather than memorizing fixed spatial patterns. Training was performed using the Adam optimizer^83^ with an initial learning rate of 1×10^−4^. To constrain model complexity and mitigate overfitting, we applied L_2_ weight decay 1×10^−4^ during optimization. The network was trained to minimize the Cross-) defined as:

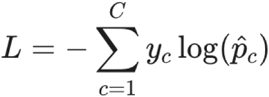

where C is the total number of classes (17), y_c_ is the binary indicator (0 or 1) of the true class label, and p_c_ is the predicted probability for class c derived from the softmax-normalized model logits.

Learning rates were managed via a MultiStepLR policy, decaying by a factor of 0.5 at epochs 10 and 20 to refine weight updates during convergence. Model parameters were initialized using Kaiming (He) uniform initialization, the default for the MONAI ResNet implementation. To ensure robust generalization, we employed an early stopping protocol^84^. Training was terminated if the class-weighted F1 score on the validation set failed to improve for 5 consecutive epochs. All experiments were implemented in Python using PyTorch^85^ and MONAI^86^ and executed on an NVIDIA RTX 6000 Ada Generation GPU.

<u>Decision-level ensemble integration of both 2D and 3D models</u>: To mitigate slice-specific artifacts (e.g., punctate noise or signal sparsity) observed in pure 3D architectures, we implemented a dual-stream decision-level ensemble framework. Single-cell inputs were independently evaluated using both the fine-tuned 2D DeepLoc model and the optimized 3D ResNet-50. The final consensus prediction, y_hat, was derived by calculating the unweighted arithmetic mean of the softmax probability vectors (P) from both component models:

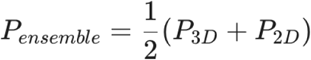

This integration leverages the complementary strengths of the two architectures: the 2D model’s projection-based invariance stabilizes predictions against volumetric noise artifacts, while the 3D model resolves complex spatial topologies that are obscured by axial projection.

## Localization maps and trajectory analysis

<u>Data filtering and age stratification:</u> The ensemble model was applied to a genome-scale dataset comprising approximately 10.7 million young cells (unbudded) and 79.4 million total aged cells across ∼5,661 yeast strains. To ensure high-confidence proteomic profiling, single-cell localization probability vectors across 17 classes—including 16 subcellular compartments and a ‘None’ category representing unexpressed cells—were subjected to quality-control filtering. Cells were excluded if the predicted probability of the ‘None’ class exceeded 0.6, indicating lack of detectable protein expression.

The remaining quality-filtered cells were stratified into six replicative age cohorts based on the quantification of WGA-stained bud scars: Young, 3–5, 6–8, 9–11, 12–14, and 15+ scars. To define a robust subcellular localization signature for each strain at every aging stage, we computed a strain-level morphological centroid by averaging the 17-dimensional probability vectors of all constituent cells within a cohort. To ensure statistical reliability, age cohorts containing fewer than 50 cells were excluded from downstream analysis, yielding a final dataset of 3,866 strains with comprehensive age-resolved profiles.

<u>Localization map:</u> To visualize the global landscape of proteome remodeling across the replicative lifespan, we employed t-Distributed Stochastic Neighbor Embedding (t-SNE). The 17-dimensional strain centroids were projected into a two-dimensional embedding space using the *scikit-learn* implementation. We utilized a cosine distance metric to capture directional similarity between localization profiles and a perplexity parameter of 30.

To facilitate direct temporal comparisons between age groups, all cohorts were plotted onto a shared global coordinate system defined by the aggregate structure of the entire dataset. This approach enables the detection of subtle shifts in protein distribution that occur as cells transition through successive replicative cycles.

<u>Trajectory analysis of protein localization trends:</u> To visualize age-dependent changes in protein localization, we generated a single shared t-SNE embedding from the 17-dimensional class probability vectors of all strains across all age cohorts. The embedding was initialized using Principal Component Analysis (PCA) on the pooled dataset to establish global consistency. All t-SNE plots were then scaled and aligned to this common embedding space, enabling direct visual comparison of localization profiles across the six age bins (Young, 3-5, 6-8, 9-11, 12-14, 15+).

Longitudinal tracking was performed by mapping the identity of the same protein from one age bin to the next (*B*_i_→*B*_i+1_). To maintain visual clarity while capturing the most prominent biological signals, we projected the trajectories of the strains representing the top 40% of these high-dimensional shifts (per organelle class and age transition) onto the 2D embedding. This visualization displays the individual trajectories of the dynamic proteins, illustrating the prevailing trends of localization change for each compartment as a function of replicative age.

Organelle-specific boundaries were defined using convex hulls. For each organelle class, the convex hull was computed as the smallest convex polygon enclosing the positions of all strains assigned to that compartment in the shared embedding. These hulls outline the core spatial region occupied by each organelle class. The trajectories and hulls together provide a direct visual summary of prevailing localization trends across aging.

## SAFE analysis

Spatially localized biological processes were identified using a customized implementation of the SAFE algorithm^22^. Strains were embedded in two dimensions by t-SNE using high-dimensional localization features. Local neighborhoods were defined as all strains within a radius set to the 1st percentile of the pairwise Euclidean distance distribution. For each GO term, enrichment within each neighborhood was evaluated using a one-tailed hypergeometric test, yielding a strain × GO-term matrix of p-values. Enrichment landscapes were calculated as −log₁₀(p) and normalized to the range [0, 1]. Significant neighborhoods were called using Benjamini–Hochberg FDR control (α = 0.05). A GO term was classified as region-specific if ≥65% of its significant neighborhoods fell within 2× the neighborhood radius. Region-specific terms were clustered by average-linkage hierarchical clustering using Jaccard distances computed from sets of significant neighborhoods. Clusters with ≥75% similarity were defined as functional regions, which were subsequently merged into meta-regions based on centroid proximity (<1.5× radius). Each strain was assigned a final color as the weighted mean of meta-region colors, using the squared normalized enrichment scores (O²) as weights, producing a continuous and interpretable functional landscape across the map.

## Organelle segmentation and protein concentration measurement

For each single-cell mNeonGreen (mNG) crop, voxels outside the cell mask were set to zero across all z-slices. Within the masked cell region, organelle-associated signal was separated from cytosolic signal using two alternative thresholding approaches applied to the full 3D stack: Yen’s threshold and a percentile threshold retaining the top 6% of voxel intensities^87,88^. For each approach, the mean intensity ratio of organelle signal to cytosolic signal was computed, and the approach yielding the larger ratio was selected for that cell.

For cells classified as cytosol by the ensemble model, any detected organelle mask and intensity were reassigned to the cytosolic compartment (i.e., organelle mask and intensity were set to 0). For cells classified as none, both organelle and cytosolic masks/intensities were set to 0.

As an additional quality-control step, the fraction of the cell occupied by the organelle mask was calculated. If the organelle mask exceeded 20% of the cell area/volume—most commonly due to minor misalignment between the mNG signal and the cell mask that introduced background and distorted thresholding—the original mNG stack was re-thresholded using a two-step procedure: (i) a top-50% intensity cutoff to remove dark background, followed by (ii) the top-6% cutoff to define organelle signal. Cells classified as vacuole or vacuolar_membrane were exempt from this re-thresholding because these compartments can legitimately occupy >20% of the cell.

Protein abundance was defined as the total mNG signal within the organelle mask, whereas protein concentration was defined as the mean mNG intensity within the organelle mask.

## Measurement of organelle size

Organelle and cytosolic masks were obtained for each cell using the selected thresholding method as described above. Cell size was defined as the sum of the organelle and cytosolic mask volumes for each cell. Strains were grouped by their predicted localization class and age. For organelle size comparisons, the plasma membrane (cell periphery), vacuolar membrane, nuclear membrane, and nucleolus were not treated as independent compartments, but instead were included with their corresponding organelles.

Representative strains for each organelle class were selected using the following criteria: (i) strong localization to the target organelle in young cells (localization score >0.8), (ii) no substantial increase in aggregation during aging, and (iii) relatively stable expression during aging. Strains labeled cytosol or none were excluded. From the remaining strains, only the top 20% by protein abundance within each organelle class were retained to prioritize well-expressed markers.

Organelle size was defined as the 3D organelle mask volume, quantified as the total number of above-threshold voxels assigned to the organelle in the z-stack (i.e., the sum of organelle-mask voxels across all slices). Cell size was defined as the total masked volume, computed as the sum of organelle-mask voxels and cytosol-mask voxels across the z-stack. For each organelle class and age group, sizes were summarized as the mean across the selected representative strains.

## KEGG enrichment

For both spatial reorganization (quantified from morphology-based image features; Figure 2) and concentration changes (quantified from fluorescence intensity; Figure 5), protein-level remodeling was tracked across aging stages. Using a 1.5-fold change cutoff, an average of 1,424 proteins with morphology changes was identified per stage-to-stage transition, along with a more stringent set of 764 proteins with high-confidence concentration changes. Functional enrichment was performed using ShinyGO v0.77^89^ with the KEGG pathway database for *Saccharomyces cerevisiae*. Optional “Pathway DB” background setting was used to limit the statistical universe to genes with KEGG annotations, which improves sensitivity for detecting coordinated changes within annotated pathways while remaining aligned with KEGG’s scope. Enrichments were considered significant at FDR < 0.05, and redundancy reduction was applied to merge overlapping terms.

## Flow network of inter-compartmental protein relocalization during aging

Proteins with robust age-dependent relocalization at the strain level (≥1.5× change in the top-1 label and ≥2× change in the secondary organelle label) and at the single-cell level (≥20% of cells showing a ≥2× change), together with clear, visually validated transitions, were included in this plot. Cytoscape was used to visualize inter-compartmental protein relocalization across age stages: node size is scaled to total outgoing flux (number of proteins exiting each organelle), and edge thickness is proportional to the number of strains relocalizing between the connected compartments.

## Protein Image-based Functional Annotation (PIFiA) for protein association analysis

<u>Detecting protein associations in young cells via adaptive thresholding:</u> We extracted 2,560 cell-level morphological features from the final feature layers of the trained deep learning models, comprising 512 features from the 2D network and 2,048 features from the 3D network. To ensure data quality, analysis was restricted to cells exhibiting a maximum class probability of at least 0.4. We subsequently applied Principal Component Analysis (PCA) to these high-confidence cells, generating 111 single-cell principal component (PC) embeddings. For downstream analysis, these PC features were aggregated by computing strain-level mean embeddings across all single-cell observations of each strain. Strains were retained for clustering only if they met strict inclusion criteria: representation by at least 50 single-cell measurements and a confident average localization probability (>0.4). This filtering ensured that only strains with robust, high-quality morphological profiles overlapping with the feature dataset were used for subsequent analysis.

To map morphological profiles to protein complexes, we compiled protein-complex membership from a curated *Saccharomyces cerevisiae* complex annotation table^90,91^, filtered to retain only genes present in the morphological dataset. Each complex’s constituent genes and associated subcellular localizations were parsed into structured lists and dictionaries to allow cross-referencing. Protein interaction network was obtained from the BioGRID v4.4.248 dataset, restricted to physical interactions. To construct the network for downstream enrichment analyses, we filtered this global dataset to retain only those interactions where both partners were present in our morphological screening library.

Morphology-defined protein clusters were identified using hierarchical clustering on strain–strain Pearson correlation matrices derived exclusively from single-cell morphological features. For each pair of strains *i* and *j*, morphological similarity was quantified using the Pearson correlation coefficient:

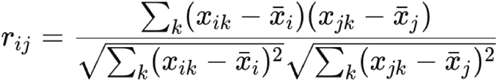

where *x_ik_* denotes the *k*^th^ morphological feature of strain *i*. Pairwise distances were defined as *d_ij_=*1 *– r_ij_*, and average-linkage clustering was applied to generate a global dendrogram. To identify robust clusters independent of a single dendrogram cut, we employed a stability-based adaptive threshold scanning strategy consistent with prior analyses of morphology-derived hierarchical feature representations^57^. The dendrogram was scanned using a step size of 0.06 over a distance range up to 0.55 (corresponding to Pearson correlation ≥0.45). Candidate clusters were required to persist across at least two consecutive thresholds, meet a minimum size of three strains, exhibit high compositional stability across cuts (overlap ≥93%), and show limited size fluctuation (±10%), yielding morphology-stable “root” clusters. Near-duplicate clusters arising from adjacent dendrogram cuts were consolidated by grouping clusters with highly overlapping membership (Jaccard index ≥0.95) and selecting a single canonical representative per group, producing a non-redundant set of stable morphology-defined clusters referred to as “canonical” clusters.

Validation of canonical clusters was performed by comparing them with curated protein complex annotations and documented protein interaction networks. For each cluster, overlap with all annotated complexes was evaluated, and the best-matching complex was identified. True positives (TP), false positives (FP), and false negatives (FN) were computed based on set overlap, from which precision, recall, and F1 scores were derived.

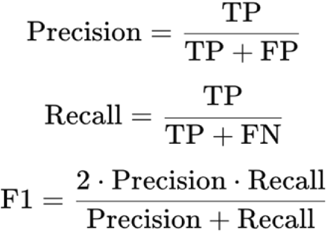

Clusters were designated as high confidence if they satisfied either of two complementary, cluster-level evidence criteria. First, clusters exhibiting substantial correspondence to curated protein complexes were retained, defined by a precision greater than 0.4, indicating that at least 40% of cluster members belonged to the same annotated complex. Second, clusters lacking a dominant complex annotation were retained if they exhibited cohesive internal interaction structure at the cluster level, defined by both (i) broad participation of proteins in protein interaction network, requiring that more than 80% of cluster members have at least one previous reported interaction with another member of the same cluster, and (ii) elevated overall within-cluster protein interaction network density (>0.25), meaning that more than 25% of all possible protein–protein pairs within the cluster are supported by experimentally observed interactions. For a cluster *C*, within-cluster protein interaction network density was computed as

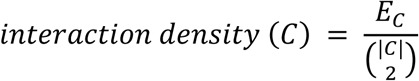

where *Ec* denotes the total number of experimentally documented interactions among cluster members, and (|*C*|, 2) represents the total number of unique protein–protein pairs that can be formed among the |*C*| proteins in the cluster. Together, these criteria ensure that protein interaction network-supported clusters are not driven by isolated interaction pairs or hub-and-spoke connectivity, but instead represent globally interconnected interaction modules. This unified criterion integrates curated complex coherence with interaction-network support while preserving morphology-derived cluster stability as the primary discovery signal.

<u>Detecting age-associated changes in protein complexes and associations</u>: To assess age-associated remodeling of protein complexes/associations, we analyzed high-confidence canonical clusters identified independently at each age bin as described above. For each protein present in high-confidence clusters at a given age, an age-specific interaction neighborhood was defined as the set of BioGRID-documented physical interaction partners that co-occur/co-cluster with that protein in the same cluster. Interaction neighborhoods were compared stepwise between consecutive age bins to classify partners as retained, decreased/lost, or increased/gained at each transition. Because only previously documented interaction partners were included in this analysis, the gain/loss of co-clustering partners across aging bins reflect increase/decrease of their PIFiA feature profile co-clustering during aging. Proteins exhibiting at least one gained or lost interaction at any transition were designated as changing proteins. By focusing on BioGRID-documented interaction partners, this analysis was not intended to discover new interactions; instead, it focused on quantifying age-associated changes in previously characterized interactions. Proteins that were present in high-confidence clusters at an earlier age but absent from all clusters at the subsequent age were classified as interaction-loss events, reflecting complete disappearance of their cluster-associated interaction neighborhoods.

Similarly, for age-associated changes in protein complex organization, we analyzed high-confidence canonical clusters using curated protein complex annotations. Analyses were performed independently for each age bin. For each protein complex represented by at least one high-confidence cluster, we computed the mean pairwise Pearson correlation among member proteins as a measure of intra-complex feature profile coherence. Absolute correlations and member counts were visualized as heatmaps. To quantify age-dependent changes relative to the young state while explicitly capturing member loss, we computed a penalized coherence ratio in which missing members were assigned a correlation of zero before normalization. This approach ensures the ratio is sensitive to both the spatial divergence of the remaining subunits and the loss of any subunit(s). Together, these analyses revealed progressive, age-associated declines and rewiring in complex-level coordination during replicative aging.

## Limited Proteolysis and TMT-Mass Spectrometry (LiP-MS) analysis

The LiP-MS was carried out as in our previous work ^59^.

<u>Sample Preparation:</u> Three biological replicates of wild-type yeast cells (BY4741) were cultured with YEPD at 30°C overnight in glass tubes on rotating drums to OD_600_ around 0.6. For each unit, about 3 OD_600_ were collected by centrifugation (3,000g, 20 seconds) as M-cells for biotinylation labeling of their cell wall. Before labeling, M-cells were washed 3 times with cold sterile phosphate-buffered saline (1xPBS, which is 10 mM; pH 8), resuspended in the same PBS, to 0.75mL. Keep cells on ice until adding biotin. Biotin solution, 4 mg/ml in cold pH 8 PBS, was added to the cells to make final concentration of 1 mg/ml, gently rock for 30 min at room temperature. Wash the cells with 1xPBS, which is 10 mM, pH 7.2, for 3 times to remove all free biotins. Then resuspend the cells in 250 ul PBS pH 7.2, added 30-50 ul BioMag beads to the remaining 250 ul cell suspension, kept on 4C rocker for 30 min. Add fresh medium to 6-well cell culture plate (6 ml per well), then put the labeled 3xOD_600_ M-cells into each well. Multiple plates were prepared for each run as the yield of old mother cells will be much less than 3 OD from each well due to cell loss during each medium changing step. Culture cells at room temperature within an orbital shaker to keep cells suspended and change medium every 4-8 hr to make sure the cell culture OD is not above 1.5 (more frequently in the first 12 hours due to fast growth). When change medium, place the 6-well plate on top of magnet bar to attract the M-cells, then get rid of the daughter cells in the old medium, and remove the magnet bar and ad fresh medium to resuspend the M-cells. After 36 hours, harvest the aged M-cells by magnet bar.

Young cells were prepared as following: By4741 cells were cultured in YPD overnight. Refresh cell to OD 600 of ∼0.2, and culture for 2-3 hours until OD 600 of ∼0.6. Collect cells by centrifugation (3,000 g, 20 sec). Add similar amount of BioMag beads as the old cell sample according to the cell pellets volume. Collected the cell-beads mix for downstream processing.

<u>Protein Digestion:</u> Cells were washed once with cold lysis buffer (50 mM Tris, pH 7.4, 150 mM NaCl, 0.5% Triton X-100, and 5% Glycerol), and resuspended in 0.5-1mL lysis buffer (depends on the OD). The cell suspensions were flash frozen as droplets (∼3mm) in pre-chilled 50 mL grinding jars in liquid nitrogen and grinded for 10 cycles with enough LN inside the Cryogrinder from SPEX samplePrep. After melting the samples, cell lysates were clarified by a 10min centrifugation at 15,000 g at 4°C to remove insoluble components. The concentration of clarified lysate was determined with BCA assay and diluted to 1.5 mg/mL with pre-chilled buffer (20mM HEPES pH 7.5, NaCl 150mM, 10mM MgCl_2_). The samples were brought to room temperature and digest with 8.5 ug Proteinase K (1:100 w/w) for 5 min at room temperature. The proteinase K-digested samples were immediately terminated by 7 M guanidine hydrochloride (GdnHCl) and boiled for 5 min.

After cooling down to room temperature, samples were reduced with 5 mM TCEP (tris(2-carboxyethyl)phosphine) and alkylated with 10 mM chloroacetamide in dark for 30 min. Samples were diluted with 0.1 M NH_4_HCO_3_ to reduce the concentration of GdnHCl to 0.5 M and digested with trypsin (Promega, 18 μg, which is 1:50 w/w) for 2hr at 37°C with 1mM CaCl_2_. Additional trypsin (12 μg, which is 1:75 w/w) was added to each sample to digest overnight. The samples were next acidified with formic acid (5%) and concentrated using Empore Solid Phase Extraction Cartridges prewashed with 1mL of methanol and 1mL of 0.1% Trifluoroacetic acid (TFA). The peptides were eluted with 80% acetonitrile containing 0.1% TFA after washing the cartridge with 1mL of 0.1% TFA. Peptides were dried using a SpeedVac concentrator and resuspended in water for peptide quantification.

<u>TMT labeling and MS Data Acquisition.</u> Frozen TMT6plex reagent vials were allowed to equilibrate to room temperature immediately before use. Each 0.8 mg vial was reconstituted in 40 μL of anhydrous acetonitrile. Based on peptide assay results, 18 μg of peptides from each sample were adjusted to a final volume of 50 μL with 100 mM triethylammonium bicarbonate (TEAB) and mixed with 20 μL of the appropriate TMT reagent. Labeling reactions were incubated for 1 hour at room temperature.

Labeling efficiency was assessed by analyzing 1 μL of each sample by LC–MS/MS using a 2-hour C18 reverse-phase gradient on an Orbitrap Eclipse Tribrid Mass Spectrometer (Thermo Fisher Scientific) equipped with a Nanospray Flex Ion Source and coupled to a Dionex UltiMate 3000 RSCLnano system. Labeling efficiency exceeded 99% for all detected peptides (data not shown). Differentially labeled samples were combined into a single tube, and the pooled sample volume was reduced to less than 10 μL using a SpeedVac concentrator.

<u>High pH reverse phase fractionation.</u> The combined TMT-labeled peptide mixture was resuspended in 300 μL of 0.1% trifluoroacetic acid (TFA). A high-pH reverse-phase fractionation cartridge (Pierce) was placed into a new 2.0 mL collection tube, and the peptide solution was loaded onto the cartridge. Following centrifugation at 3000 × g for 2 minutes, the eluate was collected as the flow-through fraction. The cartridge was then transferred to a new tube and washed with 300 μL of ddH₂O, and the eluate was collected as the wash fraction. An additional wash using 300 μL of 5% acetonitrile in 0.1% TFA was performed to remove unreacted TMT reagent.

Nine high-pH reverse-phase fractions were sequentially eluted into new collection tubes using 300 μL of 10%, 12.5%, 15%, 17.5%, 20%, 22.5%, 25%, 50%, and 80% acetonitrile in 0.1% TFA. All fractions were evaporated to dryness by vacuum centrifugation and subsequently reconstituted in buffer A (5% acetonitrile in 0.1% formic acid) prior to LC–MS analysis.

<u>Mass spectrometry data acquisition.</u> TMT-labeled peptide fractions were analyzed on an Orbitrap Eclipse Tribrid Mass Spectrometer (Thermo Fisher Scientific) equipped with a Nanospray Flex Ion Source and coupled to a Dionex UltiMate 3000 RSCLnano system. Peptides were first loaded onto an Acclaim PepMap 100 C18 trap cartridge (0.3 mm inner diameter × 5 mm length; Thermo Fisher Scientific) at a flow rate of 8 μL/min using the loading pump.

Analytical separation was performed using a 75 μm inner diameter microcapillary column packed in-house with 250 mm of 1.9 μm ReproSil-Pur C18-AQ resin (Dr. Maisch). Column temperature was maintained at 50 °C using an AgileSLEEVE column heater (Analytical Sales & Products). Mobile phase A consisted of water:acetonitrile:formic acid (95:5:0.1, v/v/v), and mobile phase B consisted of water:acetonitrile:formic acid (20:80:0.1, v/v/v). The nanoLC gradient included a 20-minute equilibration at 2% B, followed by a 5-minute ramp to 10% B, a 100-minute (or 117-minute) gradient from 10% to 30% B, a 20-minute (or 105-minute) ramp to 60% B, a 4-minute (or 55-minute) ramp to 90% B, a 10-minute wash at 90% B, a rapid return to 2% B, and a 15-minute re-equilibration period. The nanoLC flow rate was set to 0.18 μL/min.

The Orbitrap Eclipse was operated in TMT-SPS-MS3 mode. Full MS scans were acquired over a mass range of m/z 400–1600 in the Orbitrap at a resolving power of 120,000. MS2 spectra were generated by collision-induced dissociation (CID) at 35% normalized collision energy (NCE) and detected in the ion trap using turbo scan mode. Dynamic exclusion was enabled with a duration of 45 seconds. For TMT quantification, synchronous precursor selection (SPS) was applied by selecting the top 10 MS2 fragment ions, followed by higher-energy collisional dissociation (HCD) at 65% NCE and detection of MS3 spectra in the Orbitrap at a resolving power of 50,000.

<u>Database searching and data analysis.</u> LC–MS/MS data were processed using Proteome Discoverer version 3.1 (Thermo Fisher Scientific). MS/MS spectra were searched against a non-redundant Saccharomyces cerevisiae protein database downloaded from NCBI (January 12, 2022), supplemented with common contaminant proteins. Database searching was performed using SEQUEST-HT with the following parameters: no enzyme specificity, precursor mass tolerance of 10 ppm, fragment mass tolerance of 0.3 Da, and up to two missed cleavages. Carbamidomethylation of cysteine residues (+57.021 Da) and TMT labeling of lysine residues and peptide N-termini (+229.163 Da) were specified as static modifications, while oxidation of methionine (+15.995 Da) was set as a dynamic modification.

Peptide-spectrum matches were filtered to a false discovery rate (FDR) of 1% at the peptide level using Percolator within Proteome Discoverer. Reporter ion intensities were extracted from MS3 spectra, and protein quantification was performed by summing reporter ion intensities across all corresponding peptide-spectrum matches.

<u>1D profiling and differential score.</u> Peptide abundance was quantified using distributed spectral counts (dSpC). Proteinase K (PK) cleavage sites were extracted from non-tryptic peptides, and the N-terminal residue at each cleavage site was taken as the PK recognition residue (“PK residue”). For each non-tryptic peptide, its dSpC was assigned to the corresponding PK residue. For a given protein, dSpC values assigned to the same PK residue from multiple non-tryptic peptides were summed, then normalized to the total dSpC across all PK residues from that protein to estimate relative residue accessibility to PK. The resulting normalized dSpC (n-dSpC) values were plotted along the primary sequence to generate a 1D PK-accessibility profile for each protein.

To assess reproducibility across biological replicates of young cells, PK profiles for the same protein from different replicates were overlaid and a differential score was calculated by summing the absolute n-dSpC differences across all PK residues between two profiles. This differential score ranges from 0 to 200: 0 indicates identical profiles (complete overlap), whereas 200 indicates no overlap between the two 1D profiles (Figure S6E). The same approach was used to compute differential scores between young and aged cells.

Proteins exhibiting age-dependent 1D profile changes were identified by comparing, for each protein, the distribution of differential scores from young-versus-aged comparisons (9 pairs) to that from young-versus-young comparisons (3 pairs) using a one-way ANOVA. A ≥2-fold cutoff on differential scores was additionally applied to prioritize proteins with large age-associated shifts in PK-accessibility profiles.

## Complex stoichiometry

A curated set of yeast protein complexes was obtained from the *Saccharomyces cerevisiae* complex annotation table^90,91^. Protein identifiers were converted from UniProt accessions to common gene names using the Python package UniProtMapper. Complex members were filtered to retain only proteins present in our imaging screening dataset and not classified as “none” for localization; complexes with fewer than two remaining members after filtering were excluded.

For each complex and age bin, relative stoichiometric proportions were calculated from protein local concentration measurements. Specifically, the stoichiometric proportion of protein *i* in a complex was defined as:

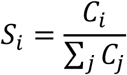

where 𝐶_𝑖_ is the mean protein concentration of member *i* and the denominator is the summed mean concentration across all retained members of the complex. An analogous proportion was computed using the standard deviation of concentration (replacing 𝐶_𝑖_ with 𝛿_𝑖_).

Age-dependent shifts in complex composition were quantified by comparing each age bin to the young reference using a custom dissimilarity metric. For each member protein, the absolute deviation from the young stoichiometric proportion was computed:

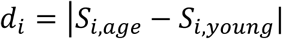

Small deviations were suppressed by setting 𝑑_𝑖_=0 when 𝑑_𝑖_< 0.15 × 𝑆_𝑖,𝑦𝑜𝑢𝑛𝑔_ (i.e., changes <15% of the young proportion). The complex-level difference was then computed as ∑_𝑖_ 𝑑_𝑖_. The same procedure was applied to the standard-deviation–based proportions. A final complex stoichiometry difference score for each complex was obtained by combining the mean-based and variability-based components (mean proportion and standard-deviation proportion) as described above.

## Protein expression heterogeneity

For each strain, single-cell heterogeneity was quantified in terms of both protein expression and morphology. Within-strain protein expression heterogeneity at each age bin was represented by the standard deviation in protein concentration across single cells at that age bin (Figure S5E). Within-strain protein morphology heterogeneity at each age bin was expressed using a 2D dispersion metric was computed in concentration space: for each strain and age bin, the centroid of the cell distribution was calculated, and the sum of Euclidean distances of individual cells to this centroid was used as an aggregate measure of dispersion (Figure S2A).

To identify strains exhibiting missed heterogeneity in local concentration—defined as substantial single-cell outliers despite a modest shift in the population mean—the fraction of cells with protein concentration below 0.67× the young-cell mean or above 1.5× the young-cell mean was calculated for each age point. Strains were classified as missed-heterogeneity candidates if the mean protein concentration at that age remained within the same 0.67×–1.5× window relative to young, yet ≥ 30% of cells fell outside that window.

## Logistic regression model for structure-based protein fate prediction

Predicted protein structures for *Saccharomyces cerevisiae* were obtained from the AlphaFold database. For each structure, we computed a high-dimensional feature set comprising 41 morphological and physicochemical descriptors (Table S6). These features quantified global compactness (e.g., radius of gyration, convex hull volume), surface composition (e.g., solvent-accessible cysteine and lysine residues), and electrostatic properties (e.g., isoelectric point), derived from structure- and sequence-based calculations.

To classify proteins based on age-associated proteomic remodeling, we implemented a binary classification framework integrating both localization dynamics and concentration changes. For each protein, organelle localization probability distributions were obtained across 17 subcellular compartments for young and aged cells. Spatial remodeling was quantified by computing the L1 distance (*D*_L1_) between young *P*_young_ and aged *P*_aged_ localization probability vectors:

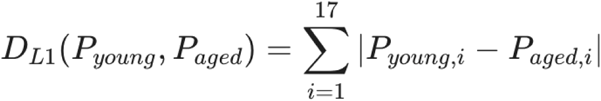

where *i* represents the probability of localization to a specific compartment (localization class). The “aging-unstable” cohort was then defined by selecting the top 30% of proteins exhibiting the highest *D*_L1_ spatial deviation, integrated with proteins exhibiting a greater than 2-fold concentration change relative to the young state. Conversely, the “aging-stable” cohort comprised the bottom 20% of proteins exhibiting minimal deviations in either metric.

Because our goal was to establish a structural feature–based model that predicts a protein’s intrinsic susceptibility to aging, we sought to distinguish proteins that are intrinsically sensitive to aging from those indirectly affected due to dependencies on age-sensitive partners. To this end, we filtered out known interacting proteins from the unstable groups, as it was unclear which member of an interacting pair acted as the driver or passenger of age-associated changes. We refined the dataset by excluding proteins heavily influenced by protein-protein interactions and removing statistical outliers (Z > 3). Recursive feature elimination reduced the input vector to 28 key features. A binary classifier was trained to distinguish aging-stable from aging-unstable proteins using this feature set. Model performance was evaluated using receiver operating characteristic (ROC) analysis on both training and held-out test datasets. In addition, we stratified the data temporally, using proteins demonstrating early-age-associated changes (age<9) for training and reserving proteins with late-age-associated changes (age>9) for independent testing. Model performance was evaluated using the same metrics as above.

Feature contributions were assessed using coefficient-based inference from the trained logistic regression model. For each feature, regression coefficients were converted to odds ratios by exponentiation, and statistical significance was evaluated using Wald tests. P-values were transformed to −log_10_(*p*) scores to enable ranking by significance. Features were ordered by p-value, and the most statistically significant predictors were visualized based on their −log_10_(*p*) values, preserving both effect size and confidence.

To evaluate evolutionary conservation of the identified biophysical determinants, the yeast-trained model was applied to a human aging proteome dataset ^60^. We focused on the 57 Ubiquitous Upregulated Proteins with Aging (UUPAs) and Ubiquitous Downregulated Proteins with Aging (UDPAs) identified in that study across at least six organs, which we treated as the aging-unstable set. As an aging-stable comparator, we curated broadly expressed housekeeping proteins reported to be proteostatically stable in the same dataset, operationalized as proteins with low age-associated variability (abundance standard deviation <0.4 across 12 tissues. For each human protein, we computed the same biophysical feature set used for yeast model training (sequence/structure-derived features as described above) and applied the pretrained model without retraining.

## Quantification and statistical analysis

Experiments, with the exception of the whole-proteome imaging screen, were repeated multiple times to confirm reproducibility. For each experiment, strains were coded with numerical identifiers, and image quantification was performed by batch processing without knowledge of strain identity. All quantifications are presented as mean ± standard error of the mean (SEM). Statistical test using Mann-Whitney U test was included in each figure and figure legend. *: p<0.05; **: p<0.01; ***: p<0.001; ****: p<0.0001.

## Supplemental information

Table S1: quantification of protein localization scores at different ages.

Table S2: KEGG enrichment for proteins with localization shift and SAFE annotations.

Table S3: list of proteins with inter-compartment localization changes at different ages.

Table S4: quantification of protein abundance and concentration changes during aging.

Table S5: PIFiA and limited proteolysis analysis of protein interaction network changes during aging.

Table S6: analysis of intrinsic biophysical and structural determinants of protein fate during aging Source Data table: number of cells quantified in each figure.

Video S1: Localization shifts of individual proteins during aging, related to Figure 2.

## Reference

1. Alberts, B., Johnson, A., Lewis, J., Raff, M., Roberts, K., and Walter, P. (2002). The Compartmentalization of Cells.

2. Lópe z-Otín, C., Blasco, M.A., Partridge, L., Serrano, M., and Kroemer, G. (2023). Hallmarks of aging: An expanding universe. Cell 186, 243–278. 10.1016/j.cell.2022.11.001.

3. Lópe z-Otín, C., Blasco, M.A., Partridge, L., Serrano, M., and Kroemer, G. (2013). The hallmarks of aging. Cell 153, 1194–1217. 10.1016/j.cell.2013.05.039.

4. Gottschling, D.E., and Nyström, T. (2017). The upsides and downsides of organelle interconnectivity. Cell 169, 24–34. 10.1016/j.cell.2017.02.030.

5. Gladyshev, V.N., Anderson, B., Barlit, H., Barré, B., Beck, S., Behrouz, B., Belsky, D.W., Chaix, A., Chamoli, M., Chen, B.H., et al. (2024). Disagreement on foundational principles of biological aging. PNAS Nexus 3, pgae499. 10.1093/pnasnexus/pgae499.

6. Zhou, W., and Cao, J. (2025). The Genomics of Aging at the Single-Cell Scale. Annu. Rev. Genomics Hum. Genet. 26, 217–243. 10.1146/annurev-genom-120523-024422.

7. Wei, Y.-N., Hu, H.-Y., Xie, G.-C., Fu, N., Ning, Z.-B., Zeng, R., and Khaitovich, P. (2015). Transcript and protein expression decoupling reveals RNA binding proteins and miRNAs as potential modulators of human aging. Genome Biol. 16, 41. 10.1186/s13059-015-0608-2.

8. Janssens, G.E., Meinema, A.C., González, J., Wolters, J.C., Schmidt, A., Guryev, V., Bischoff, R., Wit, E.C., Veenhoff, L.M., and Heinemann, M. (2015). Protein biogenesis machinery is a driver of replicative aging in yeast. eLife 4, e08527. 10.7554/eLife.08527.

9. Solyga, M., Majumdar, A., and Besse, F. (2024). Regulating translation in aging: from global to gene-specific mechanisms. EMBO Rep. 25, 5265–5276. 10.1038/s44319-024-00315-2.

10. Borner, G.H.H. (2020). Organellar Maps Through Proteomic Profiling - A Conceptual Guide. Mol. Cell. Proteomics 19, 1076–1087. 10.1074/mcp.R120.001971.

11. Thul, P.J., Åkesson, L., Wiking, M., Mahdessian, D., Geladaki, A., Ait Blal, H., Alm, T., Asplund, A., Björk, L., Breckels, L.M., et al. (2017). A subcellular map of the human proteome. Science 356. 10.1126/science.aal3321.

12. Litsios, A., Grys, B.T., Kraus, O.Z., Friesen, H., Ross, C., Masinas, M.P.D., Forster, D.T., Couvillion, M.T., Timmermann, S., Billmann, M., et al. (2024). Proteome-scale movements and compartment connectivity during the eukaryotic cell cycle. Cell 187, 1490–1507.e21. 10.1016/j.cell.2024.02.014.

13. Coody, T.K., and Hughes, A.L. (2018). Advancing the aging biology toolkit. eLife 7. 10.7554/eLife.42976.

14. Denoth Lippuner, A., Julou, T., and Barral, Y. (2014). Budding yeast as a model organism to study the effects of age. FEMS Microbiol. Rev. 38, 300–325. 10.1111/1574-6976.12060.

15. He, C., Zhou, C., and Kennedy, B.K. (2018). The yeast replicative aging model. Biochimica et Biophysica Acta (BBA)-Molecular Basis of Disease.

16. Meurer, M., Duan, Y., Sass, E., Kats, I., Herbst, K., Buchmuller, B.C., Dederer, V., Huber, F., Kirrmaier, D., Štefl, M., et al. (2018). Genome-wide C-SWAT library for high-throughput yeast genome tagging. Nat. Methods 15, 598–600. 10.1038/s41592-018-0045-8.

17. Mayr, C. (2017). Regulation by 3’-Untranslated Regions. Annu. Rev. Genet. 51, 171–194. 10.1146/annurev-genet-120116-024704.

18. Kraus, O.Z., Grys, B.T., Ba, J., Chong, Y., Frey, B.J., Boone, C., and Andrews, B.J. (2017). Automated analysis of high-content microscopy data with deep learning. Mol. Syst. Biol. 13, 924. 10.15252/msb.20177551.

19. He, K., Zhang, X., Ren, S., and Sun, J. (2015). Deep Residual Learning for Image Recognition. arXiv. 10.48550/arxiv.1512.03385.

20. Hughes, A.L., and Gottschling, D.E. (2012). An early age increase in vacuolar pH limits mitochondrial function and lifespan in yeast. Nature 492, 261–265. 10.1038/nature11654.

21. Hughes, C.E., Coody, T.K., Jeong, M.-Y., Berg, J.A., Winge, D.R., and Hughes, A.L. (2020). Cysteine Toxicity Drives Age-Related Mitochondrial Decline by Altering Iron Homeostasis. Cell 180, 296–310.e18. 10.1016/j.cell.2019.12.035.

22. Baryshnikova, A. (2016). Systematic functional annotation and visualization of biological networks. Cell Syst. 2, 412–421. 10.1016/j.cels.2016.04.014.

23. Morlot, S., Song, J., Léger-Silvestre, I., Matifas, A., Gadal, O., and Charvin, G. (2019). Excessive rDNA Transcription Drives the Disruption in Nuclear Homeostasis during Entry into Senescence in Budding Yeast. Cell Rep. 28, 408–422.e4. 10.1016/j.celrep.2019.06.032.

24. Shcheprova, Z., Baldi, S., Frei, S.B., Gonnet, G., and Barral, Y. (2008). A mechanism for asymmetric segregation of age during yeast budding. Nature 454, 728–734. 10.1038/nature07212.

25. Rempel, I.L., Crane, M.M., Thaller, D.J., Mishra, A., Jansen, D.P., Janssens, G., Popken, P., Akşit, A., Kaeberlein, M., van der Giessen, E., et al. (2019). Age-dependent deterioration of nuclear pore assembly in mitotic cells decreases transport dynamics. eLife 8. 10.7554/eLife.48186.

26. Victorelli, S., Salmonowicz, H., Chapman, J., Martini, H., Vizioli, M.G., Riley, J.S., Cloix, C., Hall-Younger, E., Machado Espindola-Netto, J., Jurk, D., et al. (2023). Apoptotic stress causes mtDNA release during senescence and drives the SASP. Nature 622, 627–636. 10.1038/s41586-023-06621-4.

27. López, G., Quezada, H., Duhne, M., González, J., Lezama, M., El-Hafidi, M., Colón, M., Martínez de la Escalera, X., Flores-Villegas, M.C., Scazzocchio, C., et al. (2015). Diversification of Paralogous α-Isopropylmalate Synthases by Modulation of Feedback Control and Hetero-Oligomerization in Saccharomyces cerevisiae. Eukaryotic Cell 14, 564–577. 10.1128/EC.00033-15.

28. Kohlhaw, G.B. (2003). Leucine biosynthesis in fungi: entering metabolism through the back door. Microbiol. Mol. Biol. Rev. 67, 1–15, table of contents. 10.1128/mmbr.67.1.1-15.2003.

29. Bonfils, G., Jaquenoud, M., Bontron, S., Ostrowicz, C., Ungermann, C., and De Virgilio, C. (2012). Leucyl-tRNA synthetase controls TORC1 via the EGO complex. Mol. Cell 46, 105–110. 10.1016/j.molcel.2012.02.009.

30. Han, J.M., Jeong, S.J., Park, M.C., Kim, G., Kwon, N.H., Kim, H.K., Ha, S.H., Ryu, S.H., and Kim, S. (2012). Leucyl-tRNA synthetase is an intracellular leucine sensor for the mTORC1-signaling pathway. Cell 149, 410–424. 10.1016/j.cell.2012.02.044.

31. Kingsbury, J.M., Sen, N.D., and Cardenas, M.E. (2015). Branched-Chain Aminotransferases Control TORC1 Signaling in Saccharomyces cerevisiae. PLoS Genet. 11, e1005714. 10.1371/journal.pgen.1005714.

32. Sato, H., and Miyakawa, I. (2004). A 22 kDa protein specific for yeast mitochondrial nucleoids is an unidentified putative ribosomal protein encoded in open reading frame YGL068W. Protoplasma 223, 175–182. 10.1007/s00709-004-0040-z.

33. Zheng, F., Yu, Z., Zhuang, J., Vannur, L., Nickols, W., Wang, Y., Li, L., Tan, S.J.O., Yoo, S., Cheng, Y., et al. (2025). Metabolic environment-driven remodeling of mitochondrial ribosomes regulates translation and biogenesis. Mol. Cell 85, 4437–4451.e11. 10.1016/j.molcel.2025.10.012.

34. Palmieri, L., Vozza, A., Hönlinger, A., Dietmeier, K., Palmisano, A., Zara, V., and Palmieri, F. (1999). The mitochondrial dicarboxylate carrier is essential for the growth of Saccharomyces cerevisiae on ethanol or acetate as the sole carbon source. Mol. Microbiol. 31, 569–577. 10.1046/j.1365-2958.1999.01197.x.

35. Payet, L.-A., Leroux, M., Willison, J.C., Kihara, A., Pelosi, L., and Pierrel, F. (2016). Mechanistic details of early steps in coenzyme Q biosynthesis pathway in yeast. Cell Chem. Biol. 23, 1241–1250. 10.1016/j.chembiol.2016.08.008.

36. Nakahara, K., Ohkuni, A., Kitamura, T., Abe, K., Naganuma, T., Ohno, Y., Zoeller, R.A., and Kihara, A. (2012). The Sjögren-Larsson syndrome gene encodes a hexadecenal dehydrogenase of the sphingosine 1-phosphate degradation pathway. Mol. Cell 46, 461–471. 10.1016/j.molcel.2012.04.033.

37. Stefely, J.A., Kwiecien, N.W., Freiberger, E.C., Richards, A.L., Jochem, A., Rush, M.J.P., Ulbrich, A., Robinson, K.P., Hutchins, P.D., Veling, M.T., et al. (2016). Mitochondrial protein functions elucidated by multi-omic mass spectrometry profiling. Nat. Biotechnol. 34, 1191–1197. 10.1038/nbt.3683.

38. Barcelos, I.P. de, and Haas, R.H. (2019). CoQ10 and Aging. Biology (Basel) 8. 10.3390/biology8020028.

39. Landl, K.M., Klösch, B., and Turnowsky, F. (1996). ERG1, encoding squalene epoxidase, is located on the right arm of chromosome VII of Saccharomyces cerevisiae. Yeast 12, 609–613. 10.1002/(SICI)1097-0061(199605)12:6%3C609::AID-YEA949%3E3.0.CO;2-B.

40. Papagiannidis, D., Bircham, P.W., Lüchtenborg, C., Pajonk, O., Ruffini, G., Brügger, B., and Schuck, S. (2021). Ice2 promotes ER membrane biogenesis in yeast by inhibiting the conserved lipin phosphatase complex. EMBO J. 40, e107958. 10.15252/embj.2021107958.

41. Stefan, C.J., Manford, A.G., Baird, D., Yamada-Hanff, J., Mao, Y., and Emr, S.D. (2011). Osh proteins regulate phosphoinositide metabolism at ER-plasma membrane contact sites. Cell 144, 389–401. 10.1016/j.cell.2010.12.034.

42. Manford, A.G., Stefan, C.J., Yuan, H.L., Macgurn, J.A., and Emr, S.D. (2012). ER-to-plasma membrane tethering proteins regulate cell signaling and ER morphology. Dev. Cell 23, 1129–1140. 10.1016/j.devcel.2012.11.004.

43. Mutlu, A.S., Duffy, J., and Wang, M.C. (2021). Lipid metabolism and lipid signals in aging and longevity. Dev. Cell 56, 1394–1407. 10.1016/j.devcel.2021.03.034.

44. Johnson, A.A., and Stolzing, A. (2019). The role of lipid metabolism in aging, lifespan regulation, and age-related disease. Aging Cell 18, e13048. 10.1111/acel.13048.

45. Hariri, H., Rogers, S., Ugrankar, R., Liu, Y.L., Feathers, J.R., and Henne, W.M. (2018). Lipid droplet biogenesis is spatially coordinated at ER-vacuole contacts under nutritional stress. EMBO Rep. 19, 57–72. 10.15252/embr.201744815.

46. Kohler, V., and Büttner, S. (2021). Remodelling of Nucleus-Vacuole Junctions During Metabolic and Proteostatic Stress. Contact (Thousand Oaks) 4, 25152564211016610. 10.1177/25152564211016608.

47. Pan, X., Roberts, P., Chen, Y., Kvam, E., Shulga, N., Huang, K., Lemmon, S., and Goldfarb, D.S. (2000). Nucleus-vacuole junctions in Saccharomyces cerevisiae are formed through the direct interaction of Vac8p with Nvj1p. Mol. Biol. Cell 11, 2445–2457. 10.1091/mbc.11.7.2445.

48. Aoyama -Ishiwatari, S., and Hirabayashi, Y. (2021). Endoplasmic Reticulum-Mitochondria Contact Sites-Emerging Intracellular Signaling Hubs. Front. Cell Dev. Biol. 9, 653828. 10.3389/fcell.2021.653828.

49. Quon, E., Sere, Y.Y., Chauhan, N., Johansen, J., Sullivan, D.P., Dittman, J.S., Rice, W.J., Chan, R.B., Di Paolo, G., Beh, C.T., et al. (2018). Endoplasmic reticulum-plasma membrane contact sites integrate sterol and phospholipid regulation. PLoS Biol. 16, e2003864. 10.1371/journal.pbio.2003864.

50. Qian, T., Li, C., He, R., Wan, C., Liu, Y., and Yu, H. (2021). Calcium-dependent and - independent lipid transfer mediated by tricalbins in yeast. J. Biol. Chem. 296, 100729. 10.1016/j.jbc.2021.100729.

51. Lanz, M.C., Zatulovskiy, E., Swaffer, M.P., Zhang, L., Ilerten, I., Zhang, S., You, D.S., Marinov, G., McAlpine, P., Elias, J.E., et al. (2022). Increasing cell size remodels the proteome and promotes senescence. Mol. Cell 82, 3255–3269.e8. 10.1016/j.molcel.2022.07.017.

52. Neurohr, G.E., Terry, R.L., Lengefeld, J., Bonney, M., Brittingham, G.P., Moretto, F., Miettinen, T.P., Vaites, L.P., Soares, L.M., Paulo, J.A., et al. (2019). Excessive cell growth causes cytoplasm dilution and contributes to senescence. Cell 176, 1083–1097.e18. 10.1016/j.cell.2019.01.018.

53. Baccolo, G., Stamerra, G., Coppola, D.P., Orlandi, I., and Vai, M. (2018). Mitochondrial metabolism and aging in yeast. Int. Rev. Cell Mol. Biol. 340, 1–33. 10.1016/bs.ircmb.2018.05.001.

54. Chan, Y.-H.M., and Marshall, W.F. (2010). Scaling properties of cell and organelle size. Organogenesis 6, 88–96. 10.4161/org.6.2.11464.

55. Cadart, C., and Heald, R. (2022). Scaling of biosynthesis and metabolism with cell size. Mol. Biol. Cell 33. 10.1091/mbc.E21-12-0627.

56. Li, Y., Jiang, Y., Paxman, J., O’Laughlin, R., Klepin, S., Zhu, Y., Pillus, L., Tsimring, L.S., Hasty, J., and Hao, N. (2020). A programmable fate decision landscape underlies single-cell aging in yeast. Science 369, 325–329. 10.1126/science.aax9552.

57. Razdaibiedina, A., Brechalov, A., Friesen, H., Mattiazzi Usaj, M., Masinas, M.P.D., Garadi Suresh, H., Wang, K., Boone, C., Ba, J., and Andrews, B. (2024). PIFiA: self-supervised approach for protein functional annotation from single-cell imaging data. Mol. Syst. Biol. 20, 521–548. 10.1038/s44320-024-00029-6.

58. Feng, Y., De Franceschi, G., Kahraman, A., Soste, M., Melnik, A., Boersema, P.J., de Laureto, P.P., Nikolaev, Y., Oliveira, A.P., and Picotti, P. (2014). Global analysis of protein structural changes in complex proteomes. Nat. Biotechnol. 32, 1036–1044. 10.1038/nbt.2999.

59. Domnauer, M., Zheng, F., Li, L., Zhang, Y., Chang, C.E., Unruh, J.R., Conkright-Fincham, J., McCroskey, S., Florens, L., Zhang, Y., et al. (2021). Proteome plasticity in response to persistent environmental change. Mol. Cell 81, 3294–3309.e12. 10.1016/j.molcel.2021.06.028.

60. Ding, Y., Zuo, Y., Zhang, B., Fan, Y., Xu, G., Cheng, Z., Ma, S., Fang, S., Tian, A., Gao, D., et al. (2025). Comprehensive human proteome profiles across a 50-year lifespan reveal aging trajectories and signatures. Cell 188, 5763–5784.e26. 10.1016/j.cell.2025.06.047.

61. Santos, A.L., and Lindner, A.B. (2017). Protein Posttranslational Modifications: Roles in Aging and Age-Related Disease. Oxid. Med. Cell. Longev. 2017, 5716409. 10.1155/2017/5716409.

62. García -Ruiz, S., Zhang, D., Gustavsson, E.K., Rocamora-Perez, G., Grant-Peters, M., Fairbrother-Browne, A., Reynolds, R.H., Brenton, J.W., Gil-Martínez, A.L., Chen, Z., et al. (2025). Splicing accuracy varies across human introns, tissues, age and disease. Nat. Commun. 16, 1068. 10.1038/s41467-024-55607-x.

63. Bhadra, M., Howell, P., Dutta, S., Heintz, C., and Mair, W.B. (2020). Alternative splicing in aging and longevity. Hum. Genet. 139, 357–369. 10.1007/s00439-019-02094-6.

64. Cesaro, L., Pinna, L.A., and Salvi, M. (2015). A Comparative Analysis and Review of lysyl Residues Affected by Posttranslational Modifications. Curr. Genomics 16, 128–138. 10.2174/1389202916666150216221038.

65. Baldensperger, T., and Glomb, M.A. (2021). Pathways of Non-enzymatic Lysine Acylation. Front. Cell Dev. Biol. 9, 664553. 10.3389/fcell.2021.664553.

66. Paulsen, C.E., and Carroll, K.S. (2013). Cysteine-mediated redox signaling: chemistry, biology, and tools for discovery. Chem. Rev. 113, 4633–4679. 10.1021/cr300163e.

67. Hess, D.T., Matsumoto, A., Kim, S.-O., Marshall, H.E., and Stamler, J.S. (2005). Protein S-nitrosylation: purview and parameters. Nat. Rev. Mol. Cell Biol. 6, 150–166. 10.1038/nrm1569.

68. Dubreuil, B., Sass, E., Nadav, Y., Heidenreich, M., Georgeson, J.M., Weill, U., Duan, Y., Meurer, M., Schuldiner, M., Knop, M., et al. (2019). YeastRGB: comparing the abundance and localization of yeast proteins across cells and libraries. Nucleic Acids Res. 47, D1245–D1249. 10.1093/nar/gky941.

69. Huh, W.-K., Falvo, J.V., Gerke, L.C., Carroll, A.S., Howson, R.W., Weissman, J.S., and O’Shea, E.K. (2003). Global analysis of protein localization in budding yeast. Nature 425, 686–691. 10.1038/nature02026.

70. Newman, J.R.S., Ghaemmaghami, S., Ihmels, J., Breslow, D.K., Noble, M., DeRisi, J.L., and Weissman, J.S. (2006). Single-cell proteomic analysis of S. cerevisiae reveals the architecture of biological noise. Nature 441, 840–846. 10.1038/nature04785.

71. Davidson, G.S., Joe, R.M., Roy, S., Meirelles, O., Allen, C.P., Wilson, M.R., Tapia, P.H., Manzanilla, E.E., Dodson, A.E., Chakraborty, S., et al. (2011). The proteomics of quiescent and nonquiescent cell differentiation in yeast stationary-phase cultures. Mol. Biol. Cell 22, 988–998. 10.1091/mbc.E10-06-0499.

72. Webb, K.J., Xu, T., Park, S.K., and Yates, J.R. (2013). Modified MuDPIT separation identified 4488 proteins in a system-wide analysis of quiescence in yeast. J. Proteome Res. 12, 2177–2184. 10.1021/pr400027m.

73. Kulak, N.A., Pichler, G., Paron, I., Nagaraj, N., and Mann, M. (2014). Minimal, encapsulated proteomic-sample processing applied to copy-number estimation in eukaryotic cells. Nat. Methods 11, 319–324. 10.1038/nmeth.2834.

74. Lu, P., Vogel, C., Wang, R., Yao, X., and Marcotte, E.M. (2007). Absolute protein expression profiling estimates the relative contributions of transcriptional and translational regulation. Nat. Biotechnol. 25, 117–124. 10.1038/nbt1270.

75. Lee, M.V., Topper, S.E., Hubler, S.L., Hose, J., Wenger, C.D., Coon, J.J., and Gasch, A.P. (2011). A dynamic model of proteome changes reveals new roles for transcript alteration in yeast. Mol. Syst. Biol. 7, 514. 10.1038/msb.2011.48.

76. Jarzab, A., Kurzawa, N., Hopf, T., Moerch, M., Zecha, J., Leijten, N., Bian, Y., Musiol, E., Maschberger, M., Stoehr, G., et al. (2020). Meltome atlas-thermal proteome stability across the tree of life. Nat. Methods 17, 495–503. 10.1038/s41592-020-0801-4.

77. Jones, D.T., and Cozzetto, D. (2015). DISOPRED3: precise disordered region predictions with annotated protein-binding activity. Bioinformatics 31, 857–863. 10.1093/bioinformatics/btu744.

78. Schindelin, J., Arganda-Carreras, I., Frise, E., Kaynig, V., Longair, M., Pietzsch, T., Preibisch, S., Rueden, C., Saalfeld, S., Schmid, B., et al. (2012). Fiji: an open-source platform for biological-image analysis. Nat. Methods 9, 676–682. 10.1038/nmeth.2019.

79. Schneider, C.A., Rasband, W.S., and Eliceiri, K.W. (2012). NIH Image to ImageJ: 25 years of image analysis. Nat. Methods 9, 671–675. 10.1038/nmeth.2089.

80. Meng, E.C., Goddard, T.D., Pettersen, E.F., Couch, G.S., Pearson, Z.J., Morris, J.H., and Ferrin, T.E. (2023). UCSF ChimeraX: Tools for structure building and analysis. Protein Sci. 32, e4792. 10.1002/pro.4792.

81. Longtine, M.S., McKenzie, A., Demarini, D.J., Shah, N.G., Wach, A., Brachat, A., Philippsen, P., and Pringle, J.R. (1998). Additional modules for versatile and economical PCR-based gene deletion and modification in Saccharomyces cerevisiae. Yeast 14, 953–961. 10.1002/(SICI)1097-0061(199807)14:10<953::AID-YEA293>3.0.CO;2-U.

82. Pachitariu, M., and Stringer, C. (2022). Cellpose 2.0: how to train your own model. Nat. Methods 19, 1634–1641. 10.1038/s41592-022-01663-4.

83. Kingma, D., and Ba, J. (2014). Adam: A Method for Stochastic Optimization. International Conference on Learning Representations.

84. Prechelt, L. (1998). Automatic early stopping using cross validation: quantifying the criteria. Neural Netw. 11, 761–767. 10.1016/S0893-6080(98)00010-0.

85. Paszke, A., Gross, S., Massa, F., Lerer, A., Bradbury, J., Chanan, G., Killeen, T., Lin, Z., Gimelshein, N., Antiga, L., et al. (2019). PyTorch: An Imperative Style, High-Performance Deep Learning Library. arXiv. 10.48550/arxiv.1912.01703.

86. Cardoso, M.J., Li, W., Brown, R., Ma, N., Kerfoot, E., Wang, Y., Murrey, B., Myronenko, A., Zhao, C., Yang, D., et al. (2022). MONAI: An open-source framework for deep learning in healthcare. arXiv. 10.48550/arxiv.2211.02701.

87. Yen, J.C., Chang, F.J., and Chang, S. (1995). A new criterion for automatic multilevel thresholding. IEEE Trans. Image Process. 4, 370–378. 10.1109/83.366472.

88. Sankur, B. (2004). Survey over image thresholding techniques and quantitative performance evaluation. J. Electron. Imaging 13, 146. 10.1117/1.1631315.

89. Ge, S.X., Jung, D., and Yao, R. (2020). ShinyGO: a graphical gene-set enrichment tool for animals and plants. Bioinformatics 36, 2628–2629. 10.1093/bioinformatics/btz931.

90. Balu, S., Huget, S., Medina Reyes, J.J., Ragueneau, E., Panneerselvam, K., Fischer, S.N., Claussen, E.R., Kourtis, S., Combe, C.W., Meldal, B.H.M., et al. (2025). Complex portal 2025: predicted human complexes and enhanced visualisation tools for the comparison of orthologous and paralogous complexes. Nucleic Acids Res. 53, D644–D650. 10.1093/nar/gkae1085.

91. Meldal, B.H.M., Pons, C., Perfetto, L., Del-Toro, N., Wong, E., Aloy, P., Hermjakob, H., Orchard, S., and Porras, P. (2021). Analysing the yeast complexome-the Complex Portal rising to the challenge. Nucleic Acids Res. 49, 3156–3167. 10.1093/nar/gkab077.

